# Pangenomics and machine learning reveal genetic variation to optimize carotenoids in sorghum grain

**DOI:** 10.64898/2026.09.22.753247

**Authors:** Linly Banda, Chloee M. McLaughlin, Avril M. Harder, Clara Cruet-Burgos, Adam L. Healey, Dave Flowers, Shannon Talley, Ada Stewart, Jane Grimwood, Geoffrey P. Morris, John T. Lovell, Davina H. Rhodes

**Author notes:** equal contribution (Linly Banda, Chloee M. McLaughlin).

## Abstract

Enhancing grain carotenoid concentration to increase yellowness and provitamin A content in sorghum (*Sorghum bicolor* [L.] Moench) is a major breeding objective across sub-Saharan Africa. Carotenoid breeding is constrained by costly post-harvest phenotyping and can be accelerated through genetic markers that capture functional variation in key carotenoid biosynthesis genes. *Zeaxanthin epoxidase* (*ZEP*) is a major gene controlling sorghum grain carotenoid accumulation, and SNP-based KASP markers facilitate selection of high-carotenoid grain, although quantitative variation remains among lines carrying the favorable allele. We hypothesized that additional functional variation within *ZEP* and other carotenoid biosynthesis pathway genes contributes to this variation and can be revealed by a pangenome-informed analysis. Using a 33-member pangenome reference, we characterized sequence and structural variation at *ZEP* and other key carotenoid biosynthesis genes and applied pangenome-derived genotyping to study marker diversity associated with carotenoid accumulation. Evaluation of *ZEP* in the pangenome reference revealed previously uncharacterized structural variation that is absent from the BTx623 primary reference genome. A pangenome-based association analysis identified genetic markers associated with the accumulation of carotenoids which were specific to particular pangenome reference members, including in RTx430 and SRN39, both yellow endosperm lines. Machine learning identified markers in *ZEP*, β*-OH*, *ZDS*, and *Z-ISO* as the most predictive for all carotenoid traits, suggesting a multi-locus genetic architecture underlies the accumulation of carotenoids in sorghum grain. By integrating sorghum pangenomic resources with machine learning, this study establishes a framework for pangenome-accelerated trait discovery and identifies new genetic targets for carotenoid biofortification in sorghum.

## INTRODUCTION

Sorghum (*Sorghum bicolor* [L.] Moench), a major staple cereal in sub-Saharan Africa (SSA), is a strategic target for provitamin A carotenoid biofortification to mitigate vitamin A deficiency (Cruet-Burgos et al., 2023, 2026; Dzakovich et al., 2023; McDowell et al., 2024; Zhao et al., 2019). Sorghum grain endosperm accumulates low concentrations of carotenoids, predominantly lutein (0.02–9.4 µg/g) and zeaxanthin (0.01–9.1 µg/g), and trace amounts (<1 µg/g) of provitamin A carotenoids (-carotene, β-carotene, and β-cryptoxanthin), the dietary precursors of vitamin A (Cruet-Burgos et al., 2026; McDowell et al., 2024). Yellow-grain genotypes, particularly those with yellow endosperm, accumulate higher carotenoid concentrations, making grain color a useful proxy for carotenoid content (McDowell et al., 2024). Grain yellowness is also commercially relevant in food and feed processing industries (Ekpa et al., 2018): carotenoid-rich yellow maize (*Zea mays*) is used in bakery applications as a natural colorant, and in poultry feed formulations to intensify the yellow-orange color of egg yolks (Ekpa et al., 2018; Flax et al., 2026; Liu et al., 2012). However, increasing heat and drought threaten maize production across SSA, exacerbating market volatility (Du & Xiong, 2024; Flax et al., 2026; Gachathi & Nzengya, 2026) and positioning sorghum, a climate-resilient C4 crop, as a promising alternative for carotenoid biofortification and industrial applications.

Conventional breeding has tripled sorghum β-carotene concentrations to 3 µg/g – representing 25% of the initial target of 12 µg/g (Cruet-Burgos et al., 2026) – demonstrating potential for genetic improvement. Further gains, however, are constrained by phenotyping: high-performance liquid chromatography (HPLC) is costly and often inaccessible to breeding programs in low-income countries, while grain yellowness lacks resolution for quantitative selection among high-carotenoid genotypes (McDowell et al., 2024). The oligogenic architecture underlying carotenoid accumulation makes marker-assisted selection (MAS) a promising breeding strategy (Cruet-Burgos et al., 2020, 2023; Fernandez et al., 2008; Halilu et al., 2016; Worzella et al., 1965), as demonstrated in maize, where favorable alleles of major carotenoid biosynthesis genes, including β*-carotene hydroxylase 1* (*crtRB1*) and *lycopene epsilon cyclase* (*LcyE*), have been deployed to increase carotenoid content (Babu et al., 2013; Harjes et al., 2008; Yan et al., 2010; Zunjare et al., 2017).

In sorghum, marker-trait associations (MTAs) have been identified within or proximal to genes involved in the precursor methylerythritol phosphate (MEP), carotenoid biosynthesis, and degradation pathways (Table 1) (Cruet-Burgos et al., 2020, 2023; Fernandez et al., 2008; McDowell et al., 2024). However, MEP and carotenoid degradation genes may be less suitable primary targets for MAS as their products support multiple essential metabolic pathways (Gupta & Hirschberg, 2022; L. Zhao et al., 2013), and selection for variation at these genes may result in flux-diverting and/or broad pleiotropic effects that do not increase carotenoid accumulation.

**Table 1.** Previously reported marker-trait associations (MTAs) for carotenoid composition and content in sorghum grain Gene ID ^†^ Enzyme Pathway Evidence type: trait.

| Gene ID<sup>†</sup> | Enzyme | Pathway | Evidence type: trait |
| --- | --- | --- | --- |
| Sobic.006G097500; *ZEP* | zeaxanthin epoxidase | Biosynthesis | GWAS: zeaxanthin, zeaxanthin/lutein ratio, total carotenoid, grain color <sup>‡</sup>(Cruet-Burgos et al., 2020; McDowell et al., 2024) |
| Sobic.004G281900; *MDS* | 2-C-methyl-d-erythritol 2,4-cyclodiphosphate synthase | MEP, precursor | GWAS: zeaxanthin (Cruet-Burgos et al., 2020) |
| Sobic.003G103300; *DXR* | deoxyxylulose reductoisomerase | Biosynthesis | GWAS: zeaxanthin (Cruet-Burgos et al., 2020) |
| Sobic.002G072400; *ZDS* | δ-carotene desaturase | Biosynthesis | GWAS: zeaxanthin (Cruet-Burgos et al., 2020) |
| Sobic.002G389000; *CYP97A* | carotene β-ring hydroxylase | Biosynthesis | GWAS: β-carotene (Cruet-Burgos et al., 2020) |
| Sobic.006G232600; *PDS* | phytoene desaturase | Biosynthesis | GWAS: β-carotene, QTL: β-carotene, zeaxanthin, total carotenoid (Cruet-Burgos et al., 2020) |
| Sobic.002G353300; *GGPPS* | geranylgeranyl diphosphate synthase | MEP, precursor | GWAS: β-carotene (Cruet-Burgos et al., 2020) |
| Sobic.003G197400; *LcyE* | lycopene epsilon cyclase | Biosynthesis | GWAS: lutein, QTL: lutein (Fernandez et al., 2008; McDowell et al., 2024) |
| Sobic.004G268500; *NCED* | 9-*cis*-epoxycarotenoid dioxygenase | Degradation | GWAS: lutein (McDowell et al., 2024) |
| Sobic.007G170300; *CCD8* | carlactone synthase | Degradation | GWAS: zeaxanthin (McDowell et al., 2024) |
| Sobic.002G225400; *CYP707A* | ABA 8'-hydroxylase | Degradation | GWAS: zeaxanthin (Cruet-Burgos et al., 2023) |
| Sobic.004G346000; *CYP97A3/LUT5* | carotene β-ring hydroxylase | Biosynthesis | GWAS: grain color (McDowell et al., 2024) |
| Sobic.006G177400; *PDS* | phytoene desaturase | Biosynthesis | GWAS: grain color ($L^*$) <sup>§</sup> (McDowell et al., 2024) |
| Sobic.002G292600; *PSY3* | phytoene synthase | Biosynthesis | QTL: endosperm color (b*), lutein, β-carotene, zeaxanthin, total carotenoid (Fernandez et al., 2008) |
| Sobic.001G509200; *CCD1* | carotenoid cleavage oxygenase | Degradation | QTL: endosperm color (b*), lutein (Fernandez et al., 2008) |
| Sobic.010G276400; *PSY1* | phytoene synthase | Biosynthesis | QTL: lutein (Fernandez et al., 2008) |
| Sobic.006G188200; *β-OH/crtRB1* | β-carotene hydroxylase | Biosynthesis | QTL: lutein (Fernandez et al., 2008) |
| Sobic.001G155300; *NCED1/Vp14* | 9-cis-epoxycarotenoid dioxygenase | Degradation | QTL: lutein (Fernandez et al., 2008) |

<sup>†</sup> Gene IDs are with respect to *S. bicolor* BTx623v5.1 genome
<sup>‡</sup> Phenotypic grain color (white, yellow kernel)
<sup>§</sup> Colorimeter based grain color, where L\* = lightness, and b\* = yellowness

Additionally, carotenoid degradation genes are often multicopy (Hao et al., 2023; Vallabhaneni et al., 2010), potentially limiting the effectiveness of selection through functional redundancy. Carotenoid biosynthesis genes, however, directly regulate the metabolic flux through the pathway and partitioning at its branch-point, making them stronger targets for MAS. Among these, *zeaxanthin epoxidase* (*ZEP*), which converts zeaxanthin to violaxanthin, has been consistently implicated in carotenoid accumulation across multiple association studies, (Cruet Burgos et al., 2020, 2023; McDowell et al., 2024), suggesting that it is a major determinant of carotenoid variation. Kompetitive Allele-Specific PCR (KASP) markers developed from loci within and near *ZEP* – notably snpSB00869 (formerly snpSB00265) – reliably distinguish high- and low-carotenoid genotypes (Cruet-Burgos et al., 2026). However, quantitative variation in carotenoid content remains among high-carotenoid genotypes carrying favorable alleles (Cruet-Burgos et al., 2020, 2023, 2026), suggesting segregation of favorable alleles at additional loci. A similar genetic architecture is observed in maize (Chander et al., 2008; Jittham et al., 2017; Kandianis et al., 2013), where favorable *LcyE* and *crtRB1* alleles account for a substantial proportion of carotenoid variation but do not consistently predict carotenoid content across genetic backgrounds (Babu et al., 2013; Diepenbrock et al., 2021; Menkir et al., 2017; Suwarno et al., 2015). We therefore hypothesized that untapped genetic variation in the sorghum carotenoid pathway contributes to this quantitative variation and could provide additional targets for accelerated breeding.

Genetic studies of carotenoid accumulation and other traits in sorghum have largely relied on the reference genome derived from BTx623 (PI564163) (Deng et al., 2024; McCormick et al., 2018; Paterson et al., 2009; Price, 2005), a genotype widely used in U.S and global breeding programs (Kane et al., 2022; Xin et al., 2021). However, single reference approaches cannot capture the full sequence and structural variation within a species, and such reference-bias may limit the identification of functional variation underlying differences in carotenoid traits. Pangenomes integrate multiple genome assemblies, enabling broader detection of SNPs/INDELs, larger structural, presence/absence, and copy number variants associated with complex trait variation (Morris et al., 2026; Ruperao et al., 2021; Schreiber et al., 2024). Approaches enabled by pangenomics have uncovered previously undetected genetic variation underlying important traits across major crops (Jiao et al., 2025; Qin et al., 2021; Song, 2026; Su et al., 2026; Yang et al., 2025). In sorghum, pangenomic studies have characterized genomic diversity of the species (Ruperao et al., 2021; Tao et al., 2021) and described loci controlling strigolactone profiles, seed shattering, dhurrin biosynthesis, and disease resistance (Maina et al., 2025; Morris et al., 2026; VanGessel et al., 2025). We therefore further hypothesized that a pangenome-enabled analysis could reveal untapped genetic variation in the sorghum carotenoid biosynthesis pathway.

In this study, we evaluated candidate genes in the sorghum biosynthesis pathway for the identification and prioritization of markers associated with differences in carotenoid traits. Building on our previous work (Cruet Burgos et al., 2020, 2023, 2026; McDowell et al., 2024), we first performed genome-wide association studies using a high-density marker set generated by variant calling against the BTx623v5.1 linear reference to identify stable and/or novel marker-trait associations. We next leveraged a 33-member sorghum pangenome reference to characterize genetic variation across *ZEP* and other key carotenoid biosynthetic candidate genes. Finally, we applied machine learning algorithms to prioritize loci within candidates across the carotenoid biosynthetic pathway – based on the observed pangenome-derived MTAs – and evaluated the contribution of the trait-associated markers to the prediction of β-carotene, lutein and zeaxanthin content. Together, this study leverages pangenomic and machine learning approaches to identify new genetic targets for sorghum carotenoid biofortification, capturing variation across the carotenoid biosynthesis pathway that has been missed by single-gene, single-reference approaches.

## RESULTS

### Marker-trait associations for carotenoids identified in genomic regions harboring a priori carotenoid biosynthesis candidate genes

We first sought to identify novel and previously reported marker-trait associations (MTAs) for carotenoid accumulation within or near *a priori* candidate genes in the carotenoid biosynthetic pathway (Table S1). Previous genome-wide association studies were conducted using variant sets of 341,514 SNPs (Cruet-Burgos et al., 2020) and 348,181 SNPs (McDowell et al., 2024) derived from genotyping-by-sequencing data aligned to the BTx623v3.1.1 reference genome. In this study, we leveraged existing carotenoid data (β-carotene, lutein, and zeaxanthin content) from a diverse panel of global sorghum germplasm, including members of the Sorghum Association Panel and the Carotenoid Panel (SAP + CAP germplasm) (Cruet-Burgos et al., 2023) alongside a new variant set generated from whole-genome re-sequencing data and called against the updated BTx623v5.1 reference genome (Morris et al., 2026). The filtered variant set included 6,735,268 SNPs and INDELs, representing a substantial increase in marker density relative to the previously available resource, and providing enhanced genomic resolution for the detection, validation and refinement of MTAs.

To account for the population structure observed for the high-carotenoid genotypes, we used two complementary association approaches: a mixed linear model with a kinship term to correct for relatedness (Figure 1A, Figure S1) and a more relaxed linear model with the first three principal components (PCs) of relatedness included as fixed effects (Figure S2). This strategy enabled the identification of associations while balancing control of population structure against the risk of overcorrection. The mixed linear model identified several significant associations (*p* < 0.05) across the genome for β-carotene (1,259), lutein (53), and zeaxanthin content (605). Only β-carotene and zeaxanthin content had significant associations within 250 kb of *a priori* carotenoid biosynthesis genes. For zeaxanthin content, eleven significant associations were detected (Table S4) within and up to 238 kb upstream of *zeaxanthin epoxidase* (*ZEP*; Sobic.006G097500), with the strongest association in the interval corresponding to a G/A SNP (SNP_Chr06_48091898) within the gene (Figure 1B, Figure S1C, S2C). This is consistent with previous observations in which the same SNP (S06_46717975: BTx623v3.1.1, snpSB00869) was the most significantly associated with variation in zeaxanthin and total carotenoid content (Cruet-Burgos et al., 2020, 2023; McDowell et al., 2024). Significant MTAs (*p* < 0.05) were detected for β-carotene content near three carotenoid biosynthesis genes (Table S5): *beta-carotene 3-hydroxylase* (β*-OH*; Sobic.001G524751), *lycopene beta-cyclase* (*Lcy*β; Sobic.007G130400), and *violaxanthin de-epoxidase* (*VDE*; Sobic.003G277400). No significant associations were detected in proximity to *a priori* carotenoid biosynthesis genes for lutein content.

**Figure 1.**
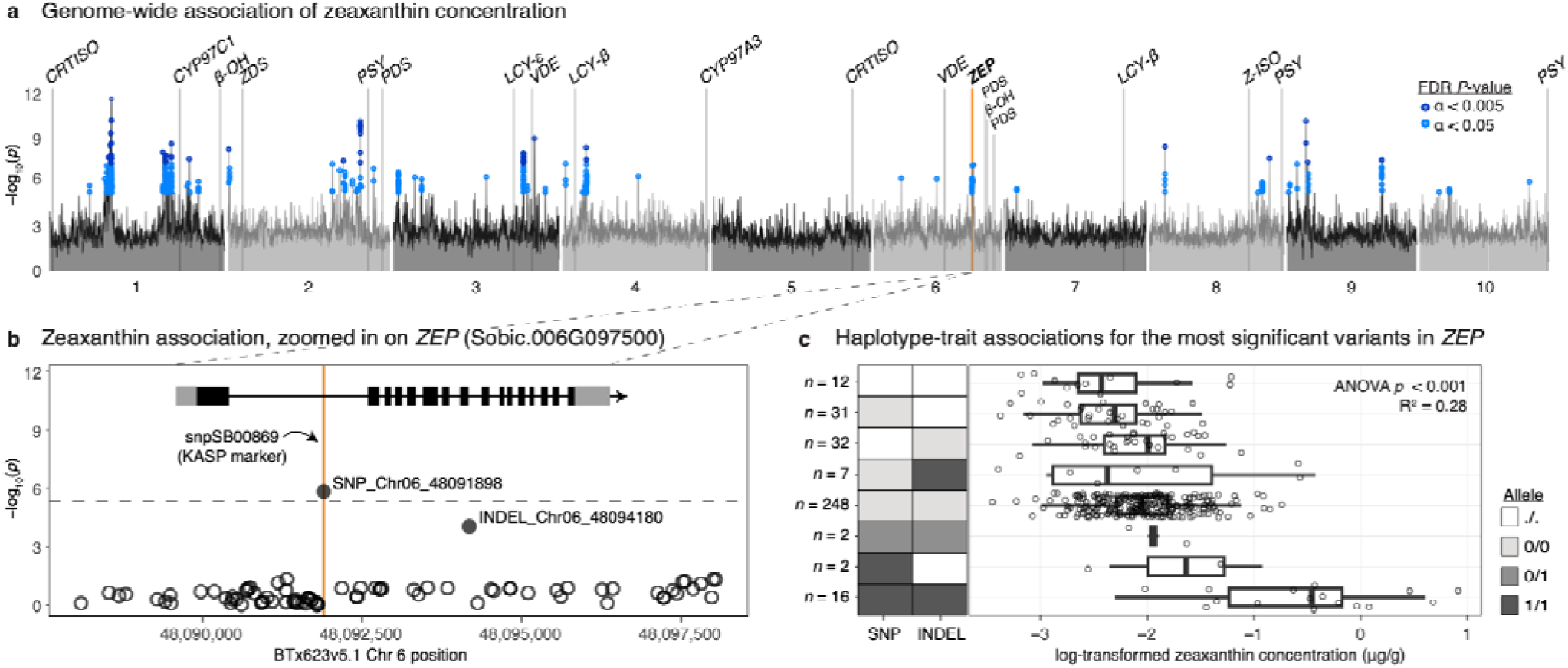
Genetic variation at the ZEP locus is associated with zeaxanthin concentration. A) Genome-wide association Manhattan plot of zeaxanthin concentration (linear mixed model accounting for kinship). The maximum -log_10_(P-value) association per 100 kb intervals is plotted genome-wide as lines. All significant associations are plotted as colored points (dark blue: FDR P-value < 0.005, light blue: FDR P-value < 0.05). Vertical lines mark the genomic position of biosynthesis candidate genes (Table S1), with the orange vertical line highlighting ZEP (Sobic.006G097500, BTx623v5.1 linear reference). B) Zoomed in association at the ZEP genomic interval ±2 kb from gene boundaries (Sobic.006G097500). Gene model features are color-coded, the light gray boxes denote 5’ (left-most) and 3’ (right-most) untranslated regions (UTRs), black boxes indicate exons. The gray dashed horizontal dashed line represents the false-discovery rate (FDR, α = 0.05) corrected threshold for genome-wide significance. The orange opaque vertical line indicates the genomic position Chr06_48091898, for which KASP marker snpSB00869 was developed to target. C) Combinations of alleles across SNP_Chr06_48091898 (SNP) and INDEL_Chr06_48094180 (INDEL) for sorghum germplasm used in association studies. Boxplots to the right of the haplotype map show distribution of log-transformed zeaxanthin concentration of phenotyped germplasm for the respective allele combinations. The number of samples of each allele combination is denoted to the left of the allele map. Multiple R² = 0.28 reports the association between zeaxanthin concentration and number of alternate alleles.

Using the linear model with the first three PCs included as fixed effects, we identified numerous significant MTAs for carotenoid traits across the genome: 76,905 for β-carotene, 137,052 for lutein, and 311,935 for zeaxanthin content. Given the model’s relaxed stringency and higher risk of false positives, we prioritized associations within *a priori* carotenoid biosynthesis genes. Significant associations for zeaxanthin content were detected within eight candidate genes (Table S6): *phytoene desaturase* (*PDS*; Sobic.002G383400), *zeta-carotene isomerase* (*Z-ISO*; Sobic.008G096800), *lycopene beta-cyclase* (*Lcy*β; Sobic.007G130400), *lycopene epsilon-cyclase* (*LcyE*; Sobic.003G197400), *cytochrome P450 monooxygenase 97C/carotenoid epsilon hydroxylase* (*CYP97C1/LUT1*; Sobic.001G308200), *beta-carotene 3-hydroxylase* (β*-OH*; Sobic.006G188200), *violaxanthin de-epoxidase* (*VDE*; Sobic.003G277400), and *ZEP.* Among these, the strongest associations were detected within *ZEP, LcyE, Lcy*β, and β*-OH* (Figure S2). For lutein content, significant associations were identified within five candidate genes (Table S7), namely *lycopene epsilon-cyclase* (*LcyE*; Sobic.003G197400), *cytochrome P450 monooxygenase 97A/carotenoid beta-ring hydroxylase* (*CYP97A3*; Sobic.004G346000), *beta-carotene 3-hydroxylase* (β*-OH*; Sobic.006G188200), *lycopene beta-cyclase* (*lcy*β; Sobic.007G130400), and *ZEP.* For β-carotene, five significant MTAs were detected (Table S8), including four SNPs within *lycopene epsilon-cyclase* (*LcyE*; Sobic.003G197400) and the G/A polymorphism in *ZEP* (SNP_Chr06_48091898).

### ZEP is a major locus controlling carotenoid content in sorghum

Given that across *a priori* carotenoid biosynthesis candidates, markers in *ZEP* exhibited the strongest association with variation in zeaxanthin content (FDR: *p* < 10^-20^ for LMM, and FDR: *p* <10^-6^ for 3PCs model) and were significantly associated with β-carotene and lutein, we performed a more detailed analysis of the genetic variation within and around this locus. In addition to SNP_Chr06_48091898, we found evidence for a previously undescribed variant approximately 2.3 kb downstream, INDEL_Chr06_48094180, to be associated with variation in zeaxanthin content (Figure 1B, Figure S2C). Using a linear model, the number of alternate alleles across these two most significant variants (SNP_Chr06_48091898 and INDEL_Chr06_48094180) explained 28% of the variation in zeaxanthin content across the SAP + CAP germplasm (Figure 1C). Variants within the *ZEP* coding sequence indicate that the interval is in a tight linkage disequilibrium block, with the notable exception of these two most significant variants, which exhibited the lowest pairwise linkage disequilibrium (*D*’ = 0.47) (Figure S3). Additionally, alternate alleles relative to the BTx623 linear reference genome across both variant sites had multiplicative effects and corresponded to the highest zeaxanthin content (Figure 1C).

### Sequence analysis across the sorghum pangenome identifies three major haplotypes at the ZEP locus

Because favorable *ZEP* alleles associated with high carotenoid content are absent from the BTx623 primary sorghum reference genome, we evaluated sequence diversity at the *ZEP* locus in 33 reference-quality *Sorghum bicolor* genome assemblies (hereafter, “pangenome members”) (Morris et al., 2026). Across pangenome members, the genomic interval containing *ZEP* has both insertions and deletions relative to BTx623 (Figure 2A), indicating additional structural diversity that may not be accurately represented in the BTx623 coordinate system. A tube map representation of the interval, in which INDEL variants ≥ 150bp were retained, demonstrates three major haplotypes across the pangenome members (Figure 2B). The predominant haplotype (*BTx623-like*) was shared by BTx623 and 21 other pangenome members mostly with white grain, including the elite African germplasm IRAT204 (PI656031), Macia (PI565121), and Mota Maradi (PI656050). A second haplotype, characterized by a 199bp deletion relative to BTx623, was observed in only four members, including one African elite accession, CSM-63 (PI655981). The third haplotype (*RTx430-like*) contained the 199bp deletion together with a 233bp insertion relative to BTx623 and was shared by seven members, including important breeding lines RTx430 (PI655996) and SRN39 (PI656027) which exhibit yellow endosperm and grain, and pale-yellow endosperm and tan grain respectively (Morris et al., 2026). Despite the minimal structural variation within the locus, we observed clear differences in sequence presence-absence (Figure 2A) and predicted gene model structure (Figure S4) across the pangenome members. Most pangenome members had one or two predicted transcript isoforms except for Wray (PI653616), which had eight.

**Figure 2.**
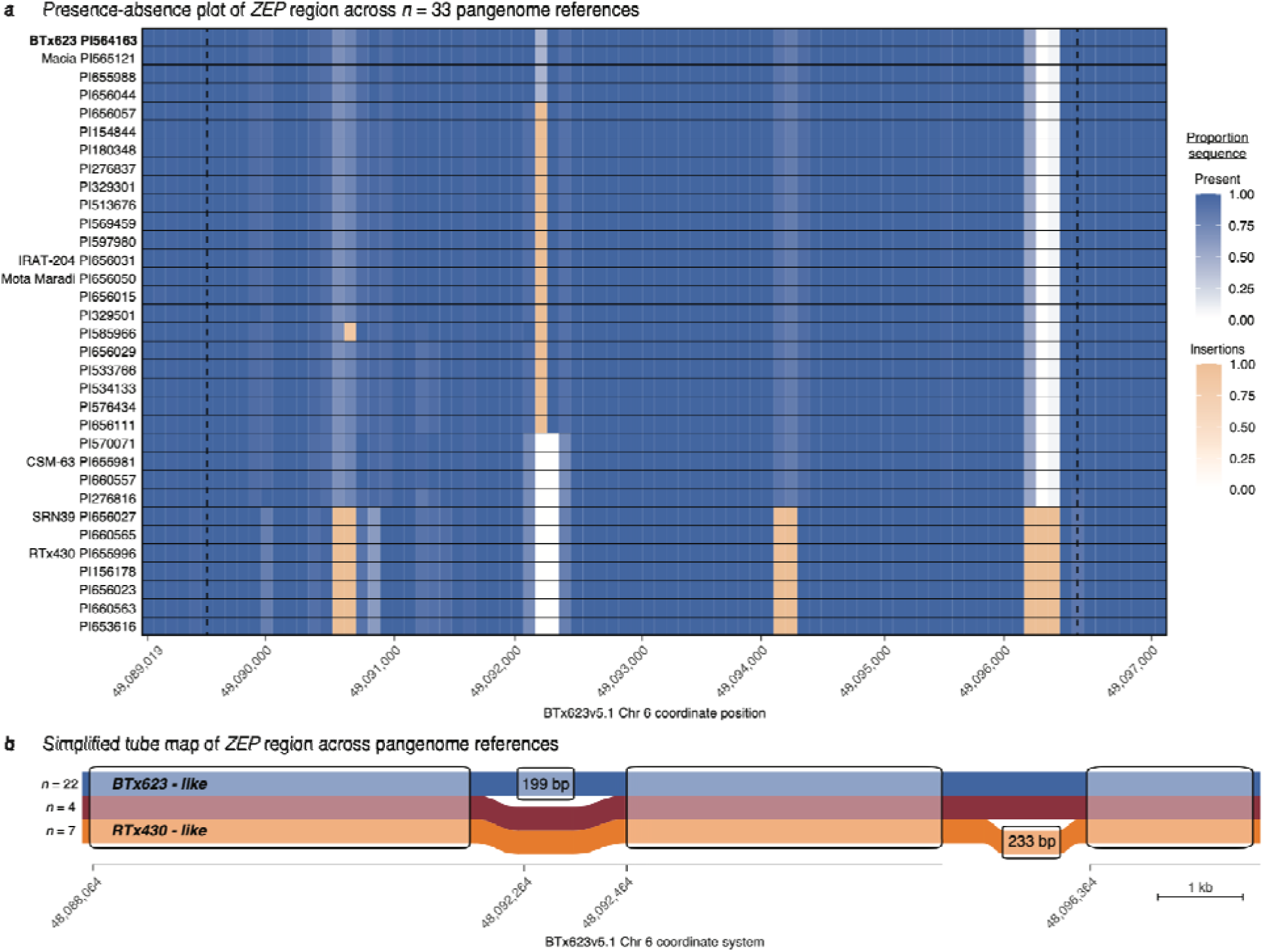
Evaluation of ZEP across sorghum pangenome members reveals structural variation not represented in the BTx623v5.1 reference genome. A) Sequence presence-absence variation in n = 33 sorghum pangenome reference member (rows) for the genomic region containing ZEP (Sobic.006G097500). The interval is plotted in a shared, expanded coordinate system to represent the sequence across all 33 genome assemblies syntenic to the primary reference BTx623v5.1 Chr06: 48,089,019 - 48,097,131. The dashed vertical black lines mark the start and end positions of the ZEP gene model as annotated in the primary reference (BTx623v5.1) within the expanded coordinate space. Each square represents a 100bp bin and is shaded according to the proportional sequence relative to BTx623 (blue: present, peach: insertion). Only insertions ≥ 10bp are indicated. B) Simplified tube map of the ZEP genomic region across n = 33 pangenome members represented in the BTx623v5.1 coordinate system. The tube map shows only large INDEL variants (≥ 150bp) across the three main ZEP haplotype paths, with the number of pangenome members belonging to each haplotype denoted on the left. Annotated snarls highlight structural features of haplotypes that deviate from the BTx623-like path, a 199bp deletion and a 233bp insertion.

### A pangenome-based genotyping approach contextualizes association studies and avoids reference bias

To further characterize the genomic region containing *ZEP* across the pangenome members, we next applied our previously described pangenome-based, reference-agnostic genotyping approach. This method utilizes the diversity of syntenic sequences across genome assemblies to identify diagnostic pangenome-derived markers (64-mers) for genotyping regions of interest (Morris et al., 2026; VanGessel et al., 2025). Using carotenoid data collected across sources (Figure S5-S6), we first calculated empirical best linear unbiased predictors (eBLUPs) of β-carotene, lutein, and zeaxanthin concentrations for *n* = 503 unique accessions (Figure S7). Broad-sense heritability estimates derived from the same linear models used to calculate the eBLUPs were lower than previously reported, with estimates of 0.44 (β-carotene), 0.66 (lutein), and 0.67 (zeaxanthin).

Of the 503 accessions, 421 unique accessions with both genotype and carotenoid phenotype data were included in the pangenome-based genotyping analysis. Here, the method was implemented as a local association study in which, for a region of interest (ROI), the association of a pangenome-derived genetic marker (64-mer) to variation in carotenoid traits is evaluated. The local pangenome-based genotyping of the *ZEP* genomic interval supported our observations from the traditional GWAS and contextualized the earlier result (Figure 3C-D).

**Figure 3.**
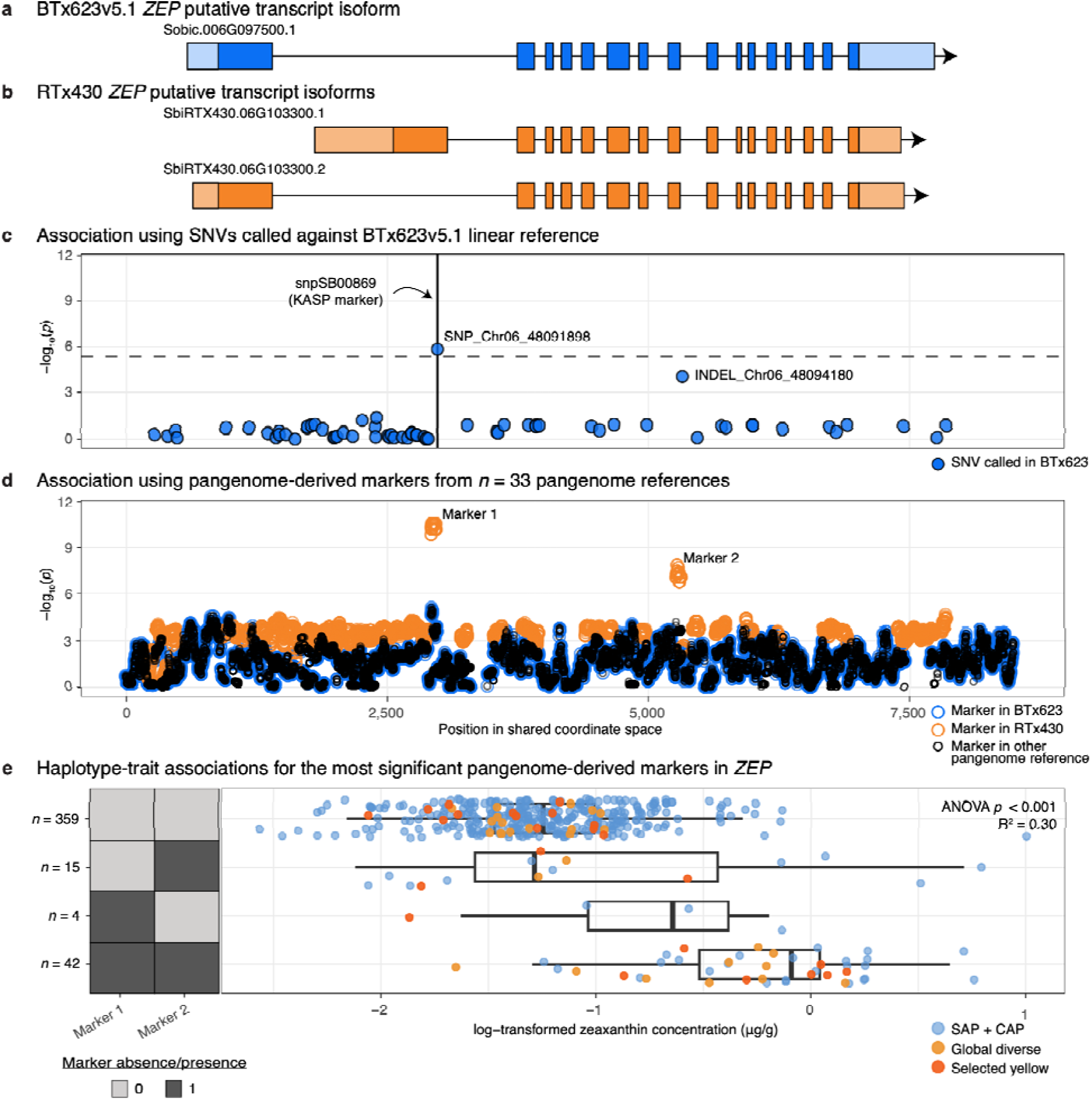
Pangenome-based associations with zeaxanthin accumulation at ZEP facilitates the identification of reference-specific favorable markers and connection to predicted transcript isoforms. A) Predicted transcript isoform for the ZEP gene model in the BTx623v5.1 genome assembly. Features of isoforms are color-coded, where light blue boxes denote 5’ (left-most) and 3’ (right-most) untranslated regions (UTRs), and dark blue boxes indicate exons. B) Predicted transcript isoforms for the ZEP gene model in the RTx430 (PI655996) genome assembly. Features of isoforms are color-coded, where light orange boxes denote 5’ (left-most) and 3’ (right-most) untranslated regions (UTRs), and dark orange boxes indicate exons. C) - log_10_(P-value) of association with zeaxanthin using variants (SNVs) called against the BTx623v5.1 primary reference. Black vertical line indicates the genomic position KASP marker snpSB00869 was developed to target in the BTx623v5.1 linear reference coordinate system. The gray dashed horizontal line is the false-discovery rate (FDR; α = 0.05) corrected threshold for genome-wide significance. Data is the same as Figure 1B, plotted in the shared coordinate system. D) -log_10_(adjusted P-value) of association with zeaxanthin using the pangenome-derived markers (64-mers). Orange points denote markers found in RTx430, blue points denote markers found in BTx623v5.1, and black points denote markers not represented in RTx430 or BTx623v5.1, but present in the other n = 31 pangenome members. All gene models and association plots (A-D) are drawn in a shared common coordinate system, based on each feature’s relative position in multiple sequence alignment space (bounds of the syntenic interval in BTx623v5.1 Chr06:48089013-48097136, bounds of the syntenic interval in the other pangenome members are reported in Table S2). E) Combinations of pangenome-derived marker absence (0, light gray) and presence (1, dark gray) across “Marker 1” and “Marker 2” annotated in (D) for sorghum germplasm used in the pangenome-based association studies (n = 421). Boxplots to the right of the marker absence/presence map show log-transformed zeaxanthin concentration of phenotyped germplasm for the respective marker combinations. The number of accessions for each combination is denoted to the left of the absence/presence map. Sorghum accessions are colored by phenotyping source: SAP + CAP germplasm (includes genotypes phenotyped in Cruet-Burgos et al. 2023 and replicated in other sources), Global diverse germplasm (novel material published in Cruet-Burgos, Banda et al. 2026), and Selected yellow germplasm (novel and previously unexplored germplasm only). One accession with an ambiguous genotype call at “Marker 1” and presence (1) at “Marker 2” of the Selected yellow germplasm group was not plotted and had a phenotype value of log(zeaxanthin concentration) =-1.71. Multiple R² = 0.30 reports the association between zeaxanthin concentration and number of present markers.

Significant associations between carotenoid variation and pangenome-derived markers of the *ZEP* genomic region existed for all carotenoid traits, with the most significant associations relating to variation in zeaxanthin content (Figure S8). When plotted in a shared coordinate space (relative position in ungapped sequence alignment of *ZEP* syntenic sequence across pangenome members), the two significant peaks of the local association using pangenome-derived markers perfectly overlap with SNP_Chr06_48091898 and INDEL_Chr06_48094180 (Figure 3C-D).

The pangenome-based association additionally revealed complexity that traditional single reference-based methods obscured. We identified two groups of pangenome-derived markers within the *ZEP* locus that are associated with zeaxanthin content. This pattern suggests that trait variation is associated with recombinant haplotypes across the locus rather than with a single causal short variant. Collectively, the pangenome-derived markers most significantly associated with zeaxanthin from each of the two clusters explained 30% of the variation in zeaxanthin content across all germplasm included in the analysis. Notably, these pangenome-derived markers most significantly associated with zeaxanthin content were specific to the RTx430 and SRN39 references, belonging to the *RTx430-like* haplotype (Figure 2B, Figure 3D). Thus, *ZEP* pangenome-derived markers associated with high zeaxanthin content are nested within the pangenome reference haplotype that is structurally most divergent from the BTx623 primary reference. Additionally, we observed annotation differences across the pangenome members: in BTx623v5.1, SNP_Chr06_48091898 is annotated as occurring in the first intronic sequence of *ZEP* (Figure 3A, C); however, examination of the orthologous gene model in the RTx430 reference genome–carrying the favorable *ZEP* alleles–reveals that the syntenic position of this SNP occurs in the first exon of the primary transcript isoform (Figure 3B, D). This highlights that relying on a single reference genome can lead to misleading functional annotation of putative causal variants.

Next, we hypothesized that the *ZEP* pangenome-derived markers most strongly associated with zeaxanthin would be enriched in a subset of yellow grain accessions selected for their favorable estimated breeding value and consequently, have higher carotenoid concentrations. Consistent with this hypothesis, we observed a higher frequency (Figure 3E) of the pangenome-derived markers (“Marker 1” and “Marker 2”) in the selected germplasm (33%) relative to the unselected germplasm (7%), indicating that favorable alleles identified in pangenome members are also present in applied breeding populations.

### Pangenome analysis of other key carotenoid biosynthesis genes (*Lcy*β, *LcyE*, and β*-OH*) identifies additional markers associated with carotenoid traits

Building on the pangenome-derived genotyping analysis of *ZEP*, we extended the approach to three additional carotenoid biosynthesis genes – *LcyE* (Sobic.003G197400), *Lcy*β (Sobic.007G130400), and β*-OH* (Sobic.006G188200) – that showed significant marker-trait associations with zeaxanthin content (*p* < 0.05) using the linear model with the relaxed population structure representation (Figure S2B-E). Of these, *LcyE* (Sobic.003G197400) and β*-OH* (Sobic.006G188200) exhibited notable pangenome-derived MTAs, while none were detected for *Lcy*β. Markers within the *LcyE* syntenic interval showed the strongest association with lutein content and were also associated with variation in zeaxanthin content (Figure S9).

When projected onto a shared coordinate system, the pangenome-derived markers from the *LcyE* interval co-localized with the association peaks identified in the linear-based GWAS, and the most significant markers were unique to the RTx430 (yellow grain) and PI513676 (white grain) reference assemblies (Figure S10, Figure S2B). Across pangenome members, the β*-OH* gene occurs in a structurally complex genomic region (Figure S11) such that when defining the reference-specific syntenic sequence to use for genotyping, approximately 35 - 52 kb of sequence was added to the bounds of the gene interval (Table S2). Although this interval expansion substantially increased the number of markers assessed in the local pangenome-based association analysis, in this instance, the broader interval captured markers distant from the target candidate gene that were associated with variation in β-carotene (Figure S12).

Interestingly, these pangenome-derived markers most associated with the β-carotene content occur in a nearby β*-1,3-glucosyltransferase* gene (Sobic.006G188600) ∼20 kb downstream from β*-OH*, and have an increased frequency in germplasm with higher accumulation of β-carotene (Figure S13). Further validation is required to confirm the nature of this signal–causal, linked to a causal locus, or simply a spurious signal.

Taken together, the pangenomic exploration of these four candidate genes (*ZEP*, *LcyE, Lcy*β, and β*-OH*) identified sequences absent from the BTx623 primary reference genome that were both poorly represented by conventional SNP/INDEL markers and significantly associated with carotenoid traits. However, this gene-by-gene approach did not fully account for the genetic variation underlying phenotypic differences or capture the additive contributions of multiple genes within the carotenoid biosynthetic pathway.

### Pangenome-based machine learning detects carotenoid associations across genes in the sorghum carotenoid biosynthetic pathway

To investigate the combined effects of variation across the carotenoid biosynthetic pathway, we extended the pangenomic analysis from individual genes to a pathway-level analysis using pangenome-derived markers from 18 *a priori* biosynthesis candidate genes (Table S1). Machine learning models (XGBoost) were developed separately for the three carotenoid traits (β-carotene, lutein, and zeaxanthin) to evaluate the predictive value of variation across the pangenome-derived markers for each trait (Figure 4A). The models identified markers that most strongly contributed to prediction accuracy, allowing the estimation of the relative importance of individual markers and candidate genes for each carotenoid trait. Of the 11,700 non-redundant pangenome-derived markers used as the initial input for model training, subsets of informative markers were identified for each trait (β-carotene, 938; lutein, 1,407; zeaxanthin, 1,490) and used to train trait-specific models. Model fit was evaluated by the R-squared of predicted versus observed values of the testing dataset, and it differed among the three traits: the zeaxanthin model had the best fit (*R*² = 0.30), followed by the lutein (*R*² = 0.17), and β-carotene (*R*² = 0.15) models (Figure 4B, Figure S14). Interestingly, this estimation of model fit across the three carotenoid traits followed the earlier trend we observed in broad-sense heritability estimates, in which variation in zeaxanthin and lutein was more attributable to genetic differences captured by pangenome-derived markers of biosynthesis candidate genes than variation in β-carotene.

**Figure 4.**
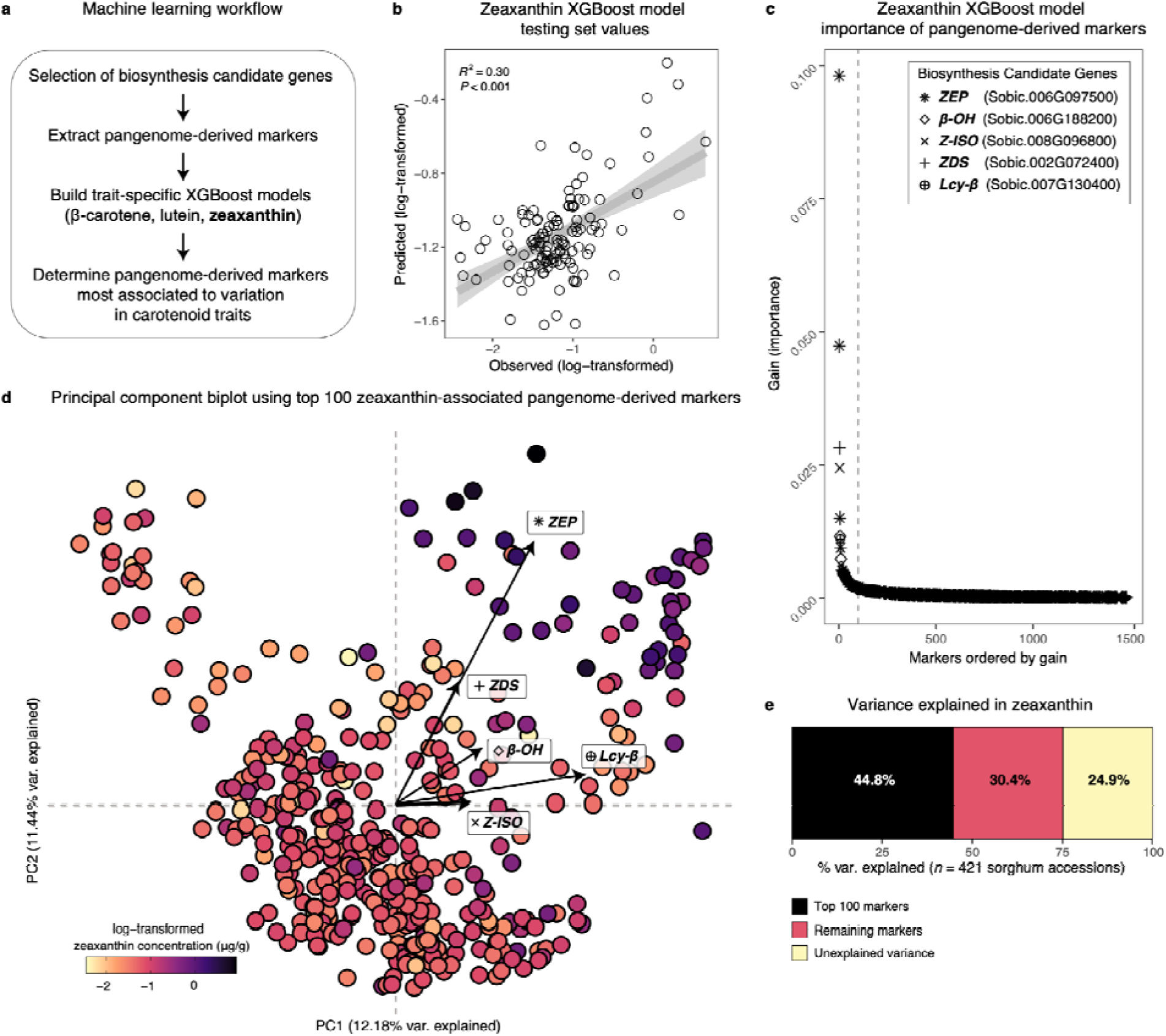
Sorghum germplasm with the highest zeaxanthin content harbor similar pangenome-derived markers across top candidate biosynthesis genes. A) Workflow for applying a machine learning approach using pangenome-derived markers. B) For the zeaxanthin XGBoost model, predicted versus observed trait values for the n = 127 accessions of the testing (hold out) data not used to train XGBoost models. C) The 1,490 pangenome-derived markers selected for training the XGBoost model for zeaxanthin prediction, ordered by gain (feature importance measure). Each point represents a marker and is styled (shape, color) according to the carotenoid biosynthesis candidate gene to which it belongs. Markers from the five candidate genes with the highest aggregated gain are additionally highlighted with a distinct point shape. In the legend, the top five biosynthesis candidate genes are ordered by the aggregated gain of markers, where the first gene reported accounts for the highest gain in model fit. The vertical dashed line indicates the 100th most informative pangenome-derived marker based on gain; markers to the left of this threshold were used to construct the biplot shown in (D). D) Principal component biplot of n = 421 phenotyped germplasm based on the 100 most informative pangenome-derived markers from the zeaxanthin XGBoost model. Points are phenotyped accessions and colored by log-transformed zeaxanthin concentration. Arrows indicate the average loading vector for the top five candidate genes by aggregated gain and have the same point styling as in (C). Vector loadings for the remaining thirteen biosynthesis candidate genes are reported in Table S3. E) Variance explained in zeaxanthin concentration by sets of pangenome-derived markers. Stacked bar chart shows the proportion of phenotypic variance in zeaxanthin across n = 421 germplasm explained by the top 100 XGBoost-selected markers (black), the 1,390 remaining markers (salmon), and unexplained variance (yellow). B-E) Figures generated using the zeaxanthin model outputs.

For each trait-specific model, we ranked the pangenome-derived markers by their contribution to model fit (gain), providing a measure of their importance for trait prediction (Figure 4C, Figure S15-S16). For the β-carotene model, the highest-ranking markers belonged to the candidate genes *VDE* (Sobic.006G049200), *ZEP* (Sobic.006G097500), *ZDS* (Sobic.002G072400), *Z-ISO* (Sobic.008G096800), *lcy*β (Sobic.004G074000) (Figure S15). For lutein, markers belonging to *ZEP* (Sobic.006G097500), *lcyE* (Sobic.003G197400), *PSY* (Sobic.008G180800), *ZDS* (Sobic.002G072400), and *PDS* (Sobic.006G177400) were the most important (Figure S16). Markers belonging to *ZEP* (Sobic.006G097500), *ZDS* (Sobic.002G072400), *Z-ISO* (Sobic.008G096800), β*-OH* (Sobic.006G188200), and *lcy*β (Sobic.007G130400) were the most informative for describing differences in zeaxanthin (Figure 4C). Two pangenome-derived markers in *ZEP* (Sobic.006G097500) showed the highest importance in the zeaxanthin model (Figure 4C) and they correspond to the two most significant peaks identified in the pangenome-based association of *ZEP* and zeaxanthin content (Figure 3D).

Notably, these same *ZEP* markers ranked within the top three most important features in both the β-carotene and lutein models (Figure S15-S16), further supporting the importance of the *ZEP* locus across carotenoid traits in sorghum.

Sorghum germplasm with the highest zeaxanthin content formed a distinct cluster in principal component space, based on the top 100 pangenome-derived markers with the highest gain (Figure 4D), accounting for 53.45% of the total model gain and 44.8% of the variance explained in zeaxanthin concentration (Figure 4E). This separation suggests that high-zeaxanthin accessions harbor unique combinations of markers at candidate carotenoid biosynthesis loci and demonstrates that the XGBoost model captured meaningful genetic structure underlying variation in zeaxanthin concentration.

We next assessed the relative contribution of pangenome-derived marker diversity at each carotenoid biosynthesis candidate gene to variation in the carotenoid traits (Supplementary File 3). For each trait-specific model and candidate gene, we summed the gain (feature importance) of pangenome-derived markers selected by the model. This aggregated gain value provides an estimate of each candidate gene’s contribution to phenotypic differences in carotenoid traits, as represented by the trait-specific XGBoost models (Figure 5A, Table S3).

**Figure 5.**
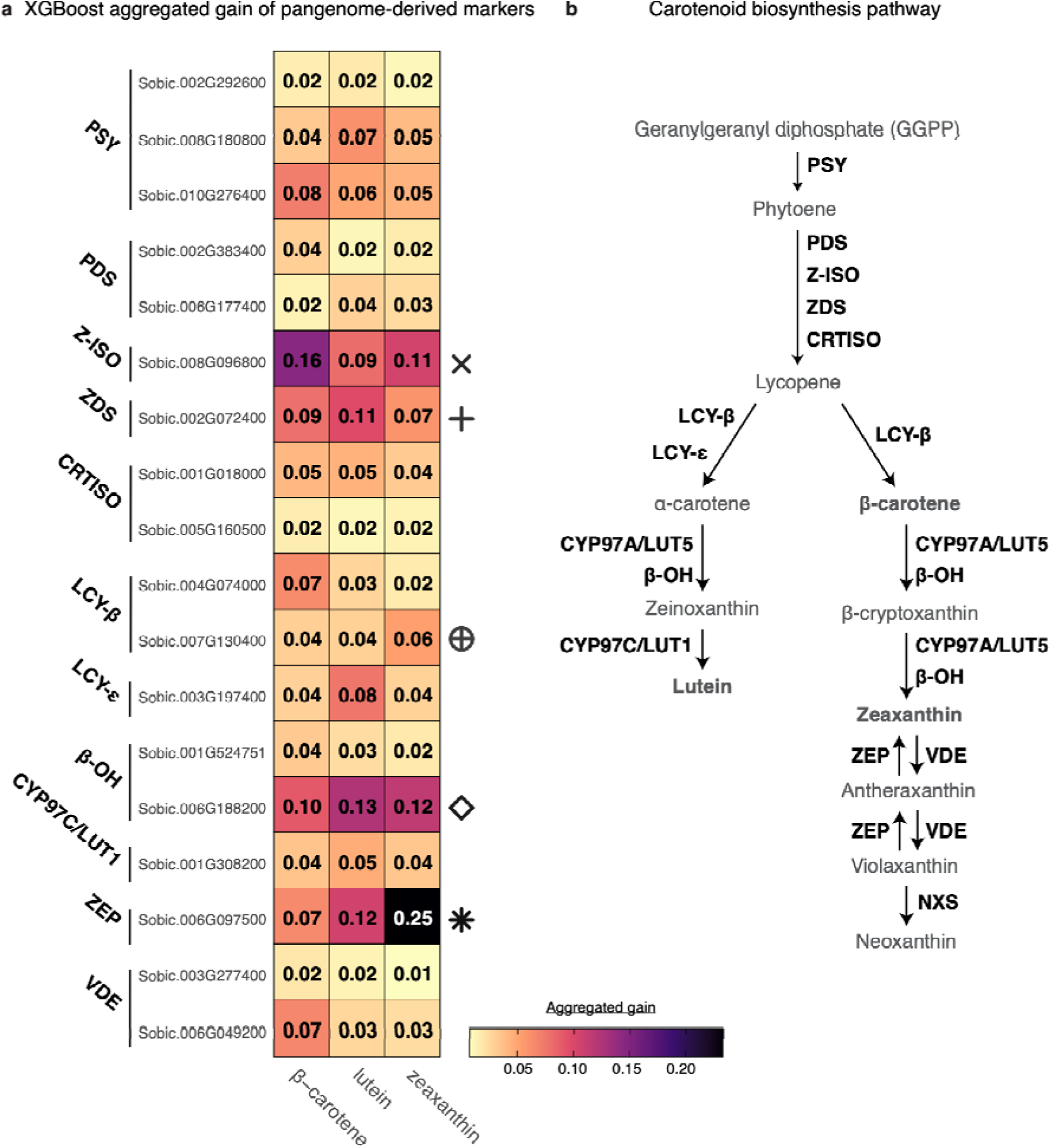
Genetic differences in pangenome-derived markers across carotenoid biosynthesis candidate genes relate to variation in carotenoid traits. A) For each candidate biosynthesis gene (Sobic IDs), trait-specific measures of gain (feature importance) aggregated across pangenome-derived markers visualized as a tileplot. Aggregated gain was calculated from XGBoost models, built separately for each trait, and represents the summed value across all pangenome-derived markers for each candidate biosynthesis gene. Multiple Sobic IDs for the same gene (PSY, PDS, CRTISO, Lcyβ, β-OH, VDE) denote paralogous genes. Fill color indicates the relative gain of pangenome-derived markers. The five candidate genes with the highest aggregated gain from the zeaxanthin model are additionally highlighted with a distinct point shape, same as in Figure 4C. B) The carotenoid biosynthesis pathway

The relative aggregated gain from the pangenome-derived markers of the candidate genes differed among carotenoid traits, ranging from 1.88% to 15.55% for β-carotene, 1.63% to 13.46% for lutein, and 1.33% to 24.69% for zeaxanthin. Variation in zeaxanthin content was again most strongly associated with variation in the pangenome-derived markers of *ZEP* and showed the strongest signal across all traits and candidate regions examined, with an aggregated gain of 24.69% (Figure 5A), followed by the markers from β*-OH* (Sobic.006G188200) and *Z-ISO*. Variation in lutein content was most associated with the markers from β*-OH* (Sobic.006G188200), *ZEP*, and *ZDS*, while that of β-carotene was most associated with the markers from *Z-ISO,* β*-OH* (Sobic.006G188200), and *ZDS*.

## DISCUSSION

### Pangenomes capture additional variation beyond linear reference-based association studies

Identifying the genetic basis of complex traits remains a central challenge in crop improvement, and pangenomic approaches offer a promising path forward by capturing genetic variation that single-reference genomes miss. Early plant pangenome studies identified variable genes associated with flowering time, maturity, and seed composition in soybean (*Glycine max*) (Y. Li et al., 2014), disease-resistance in arabidopsis (*A. thaliana*) (W.-B. Jiao & Schneeberger, 2020), and flavor in tomato (*Solanum lycopersicum*) (Gao et al., 2019), demonstrating their benefit for capturing genetic diversity across multiple genome assemblies and for revealing trait-associated differences in genotypes. Sorghum pangenomic resources now provide similar opportunities to characterize genomic diversity underlying agronomically important traits (Maina et al., 2025; S. R. Marla et al., 2025; Ruperao et al., 2021; Tao et al., 2021; VanGessel et al., 2025).

Here, we investigated genetic variation underlying carotenoid accumulation, an oligogenic trait governed by an interconnected metabolic network. We demonstrated that linear references remain effective for broad trait discovery when improved reference quality enables more comprehensive variant representation. This was evidenced by the identification of MTAs for carotenoid traits (β-carotene, lutein, and zeaxanthin) near carotenoid biosynthesis candidate genes (Figure S1A, Figure S2A), including new associations within/near *Lcy*β and *VDE,* and a 10bp INDEL (INDEL_Chr06_48094180) in *ZEP*, not previously implicated in carotenoid accumulation (Figure 1B-C). The overlap between MTAs identified by the single-reference GWAS and the pangenome-derived association for *ZEP* (Figure 3C-D), *LcyE* (Figure S10, Figure S2B) and β*-OH* (Figure S12) further confirmed that the linear-reference GWAS detected real association signals, while the lack of agreement at the *Lcy*β loci may reflect differences in marker representation, in which sequence variation within the haplotype can split the associated allele into multiple lower frequency markers, thereby reducing statistical power to detect association (Roberts et al., 2025; Voichek & Weigel, 2020). Despite the utility of the linear reference, we further demonstrated that for candidate regions involved in the accumulation of carotenoids, the sorghum 33-member pangenome reference captures and contextualizes additional sequence diversity associated with favorable trait variation.

At the *ZEP* locus, a major determinant of sorghum zeaxanthin content and carotenoid accumulation, pangenome members revealed additional sequence diversity surrounding the lead SNP (snpSB00869) (Figure 2, Figure 3), suggesting that it tags a broader functional haplotype comprising multiple linked variants within the locus. Although linkage disequilibrium may allow this SNP to capture most of the local variation, recombination or allelic heterogeneity can decouple causal variants from lead markers, limiting its ability to resolve phenotypic variation (Cobb et al., 2018; S. Marla et al., 2023; Zhou et al., 2022). Supporting this interpretation, the newly identified INDEL exhibited low linkage disequilibrium with snpSB00869 (Figure S3), and accessions carrying the favorable alleles at both loci accumulated significantly higher zeaxanthin content than those carrying only one (Figure 3E). Taken together, these findings suggest that these two variants capture partially independent loci underlying zeaxanthin accumulation and explain both the effectiveness of snpSB00869 for selecting high-carotenoid germplasm, and its limited ability to distinguish them. Consequently, selection at both loci, accounting for surrounding haplotypes, could improve genetic gain for zeaxanthin and total carotenoid content.

### Pangenome analysis reveals reference bias potentially obscuring carotenoid trait discovery

Reliance on a single linear reference genome can introduce reference bias, whereby reads carrying reference-like sequences are preferentially mapped and retained, while reads containing divergent alleles or sequences absent from the reference map poorly or remain unmapped. This can underestimate genetic diversity, particularly structural and presence-absence variation, potentially obscuring functional variants, constraining trait discovery, and contributing to unexplained phenotypic variation (Ahmad et al., 2025; Brandt et al., n.d.; Dolenz et al., 2024; Günther & Nettelblad, 2019; Sirén et al., 2021; Stevenson et al., 2013).

In this study, investigation of *ZEP* sequence diversity across the 33-member pangenome reference identified three major haplotypes, with the favorable haplotype (*RTx430-like*) differing from the BTx623 reference (Figure 2), highlighting that reliance on BTx623 alone may limit the discovery of genetic variation underlying carotenoid traits. Pangenomic analysis of the *RMES1* locus similarly revealed absence of aphid-resistance alleles from BTx623 due to a large deletion affecting NLR copy number variation within the locus (VanGessel et al., 2025). The recovery of these non-reference haplotypes highlights the importance of pangenome resources for identifying favorable alleles that would otherwise remain inaccessible using conventional, reference-based analyses. The absence of the favorable *ZEP* haplotype from pangenome reference members representing elite breeding material (Figure 2) but enrichment of favorable pangenome-derived markers in some yellow breeding lines (Figure 3E) identifies an opportunity to introgress high-carotenoid alleles into breeding populations through MAS.

It was also apparent that reference bias extends beyond variant discovery to gene annotation: the lead carotenoid-associated *ZEP* SNP (snpSB00869), annotated as intronic in the BTx623 model, occurs in the first exon of RTx430 and the other genomes carrying the favorable haplotype (Figure 3, Figure S4). This annotation shift substantially alters the variant’s predicted functional significance. Although the favorable allele may affect transcript processing possibly through altered splicing, functional validation is required. More broadly, these findings contribute to the growing recognition that reference-specific annotation artifacts can often obscure biological interpretation (Brůna et al., 2026) and further demonstrate how pangenome-based analyses provide a more comprehensive representation of functional genetic diversity than approaches that rely on a single reference genome (Lovell et al., 2026).

### Pathway-level machine learning analysis captures wider genetic variation in high-carotenoid germplasm

Metabolite accumulation reflects the coordinated activity of multiple genes within a biosynthetic pathway and is therefore unlikely to be fully explained by individual loci. While gene-level analyses resolve allelic variation within specific candidates, pathway-level approaches integrate variation across functionally connected genes, capturing cumulative additive or moderate genetic effects that may be overlooked when loci are considered independently. This approach can improve the prediction of complex phenotypes (Turner-Hissong et al., 2020), and prioritization of genomic regions associated with trait variation. We applied a pathway-level machine learning approach using pangenome-derived markers to quantify the contribution of genetic variation across carotenoid biosynthesis genes to carotenoid traits.

Among the three carotenoid traits included in this study, zeaxanthin showed the strongest predictive performance (Figure 4B, Figure S14), suggesting that its variation is more effectively captured by the pangenome-derived markers within the carotenoid biosynthesis genes. This is consistent with our previous finding that five of eleven KASP markers predicted zeaxanthin content across pre-breeding and diverse global germplasm, in contrast with only two for β-carotene and lutein (Cruet-Burgos et al., 2026). Additionally, the higher broad-sense heritability estimated for zeaxanthin compared with β-carotene and lutein, is consistent with trends reported in previous studies in sorghum and maize (Cruet-Burgos et al., 2020; Cruet-Burgos & Rhodes, 2023; Diepenbrock et al., 2021; Fernandez et al., 2008; Halilu et al., 2016; Kandianis et al., 2013; Wong et al., 2004). These findings suggest that a greater proportion of zeaxanthin variation is genetically determined and captured by genetic variation within the carotenoid biosynthesis pathway. Together with our previous observation of predominantly additive effects across contributing loci, these findings support zeaxanthin as a tractable target for MAS (Cruet-Burgos et al., 2026). The lower heritability and predictive performance of β-carotene may reflect greater environmental influence (Falconer & MacKay, 1996; Halilu et al., 2016) or genetic variation outside the candidate genes included in our model, including genes involved in precursor or degradation pathways, and other regulatory mechanisms (Cruet-Burgos & Rhodes, 2023; Diepenbrock et al., 2021; Yin et al., 2024).

Based on the relative contributions of their pangenome-derived markers in our XGBoost models, *ZEP,* β*-OH, ZDS,* and *Z-ISO* consistently ranked among the most important genes for all carotenoid traits, while markers from *Lcy*β contributed to the prediction of β-carotene and zeaxanthin, and those from *LcyE* contributed to lutein content (Figure 5). The importance of *ZDS* and *Z-ISO* likely reflects their upstream pathway positions (Figure 5) and their influence on the metabolic flux towards downstream carotenoids. *ZDS* and *Z-ISO* have been associated with carotenoid variation in cereals (Chander et al., 2008; Y. Chen et al., 2010; Wong et al., 2004; Wurtzel et al., 2012), and *ZDS* markers allow reliable selection for grain yellowness and carotenoid content in wheat (C. Dong et al., 2012; Gaponov et al., 2026). Furthermore, both are single-copy genes in maize and sorghum (Matthews, 2003), whose enzymatic functions cannot be compensated for by alternative enzymes. Together, their upstream pathway positions and lack of functional redundancy make them critical regulators of carotenoid accumulation, likely contributing to their consistent association with all carotenoid traits.

The importance of the β*-OH*-derived markers in predicting the three carotenoid traits is consistent with its role in both the α- and β-pathways (Figure 5) in which it hydroxylates the carotenes into xanthophylls. KASP markers tagging this gene–snpSB00279 and snpSB00280– accurately predict β-carotene, lutein, and zeaxanthin content (Cruet-Burgos et al., 2026). However, the most predictive β*-OH*-derived markers were in a nearby β-D-glucan biosynthesis gene, β*-1,3-glucosyltransferase* (Sobic.006G188600) (Figure S11-S13) and this may reflect linkage disequilibrium between the marker and uncharacterized regulatory or structural variants of β*-OH*, although an independent role of the β*-1,3-glucosyltransferase* cannot be excluded. We have previously identified a QTL for β-cryptoxanthin nested within a related pectin biosynthesis gene, *Galacturonosyltransferase-like 4* (Sobic.002G398400) (McDowell, 2024). These genes may influence carotenoid content through xanthophyll modification (X. Chen et al., 2021; Göttl et al., 2024) or indirectly through plastid biogenesis or grain filling (N.-Q. Dong et al., 2020; Egelund et al., 2010; Mohnen, 2008). Further association studies across the entire local haplotype containing the β*-OH* and β*-1,3-glucosyltransferase* region, coupled with linkage disequilibrium and gene expression analysis might help resolve the nature of this signal. The importance of *Lcy*β and *LcyE* to specific carotenoid traits also reflects their key branch-point positions within the pathway (Figure 5). *Lcy*β converts lycopene to β-carotene, which is subsequently converted to zeaxanthin, while *LcyE* acts in concert with *Lcy*β to produce α-carotene with subsequent modification to lutein (Cunningham et al., 1996; Harjes et al., 2008; Zeng et al., 2015). The intricate balance between the activities of these two enzymes dictates the metabolic flux partitioning into either the α- or β-branch, making them important targets for qualitative control of carotenoids.

Overall, selection of favorable haplotypes–associated with high carotenoid content– at *ZEP,* β*-OH, ZDS,* and *Z-ISO*, supports the first phase of our previously proposed breeding strategy ‘Max_Car’ aimed at maximizing total carotenoid content by increasing overall metabolic flux through the pathway (Cruet-Burgos et al., 2026). Simultaneous selection of *Lcy*β alleles associated with high β-carotene, and *LcyE* alleles associated with low lutein would support the second phase of our proposed breeding strategy, ‘Max_Beta’ which aims to increase provitamin A carotenoids, mainly β-carotene.

## CONCLUSIONS

This study demonstrates the value of integrating pangenomics and machine learning to resolve genetic variation underlying carotenoid accumulation in sorghum. Association mapping using an updated linear reference identified a *ZEP* SNP (snpSB00869) and INDEL (INDEL_Chr06_48094180) as the strongest associations with zeaxanthin content (Figure 1).

Exploration of the *ZEP* locus across pangenome references revealed three major haplotypes (Figure 2) and showed that the BTx623 reference lacks the favorable haplotype, potentially limiting its use in carotenoid trait discovery. Association mapping with pangenome-derived markers further supported these findings, with markers harboring the two *ZEP* variants showing the strongest associations with carotenoid content, and their enrichment in high-carotenoid, yellow grain accessions (Figure 3). Machine-learning identified pangenome-derived markers from *ZEP,* β*-OH, ZDS,* and *Z-ISO* as the main predictors of β-carotene, lutein, and zeaxanthin, with markers from *Lcy*β and *LcyE* additionally contributing to β-carotene/zeaxanthin, and lutein respectively (Figure 4, Figure 5). Thus, selection of favorable *ZEP,* β*-OH, ZDS,* and *Z-ISO* haplotypes could support an overall increase in carotenoid content through our previously suggested ‘Max_Car’ breeding strategy for maximising total carotenoid content, while selection for favorable *Lcy*β and *LcyE* alleles could support qualitative control of carotenoids, and contribute towards the ‘Max_Beta’ strategy for maximising β-carotene. Together, these findings provide an opportunity to accelerate molecular breeding for high carotenoid, yellow sorghum grain.

## METHODS

### Assembly of an *a priori* list of carotenoid biosynthesis candidate genes

Candidate genes associated with carotenoid accumulation in sorghum have previously been identified through homology-based annotation using maize (*Zea mays*) as a reference (Cruet-Burgos et al., 2020, 2023). The genes have been broadly categorized into the methylerythritol phosphate (MEP) precursor pathway, the carotenoid biosynthesis pathway, and the carotenoid degradation pathway. In the present study, we focused exclusively on twenty (20) genes within the carotenoid biosynthesis pathway (Table S1).

### Genome-wide association study using a denser marker set to an updated sorghum reference genome

To determine if increased marker density across the genome improved the resolution of previously identified marker-trait associations (MTAs) and facilitated the discovery of additional associated loci, we conducted a genome-wide association study (GWAS) for carotenoid traits using existing phenotypic data for β-carotene, lutein, and zeaxanthin concentrations from *n* = 350 accessions. These included lines from the sorghum association panel (SAP) and carotenoid panel (CAP), the latter comprising accessions selected based on yellow endosperm and/or yellow grain as described in (Cruet-Burgos et al., 2020, 2023). This phenotype data set is hereafter referred to as the SAP + CAP. Association analyses were performed using a recently published variant dataset filtered for 6,735,268 SNPs and INDELs (MAF > 0.05; missingness < 50%) mapped to the updated BTx623v5.1 reference genome (Morris et al., 2026). GWAS was performed in GEMMA (v.0.98.30) with default parameters. Sorghum germplasm with high carotenoid content, particularly within this previously phenotyped panel, exhibits limited genetic diversity and strong population structure, which may inflate spurious associations and complicate the identification of true MTAs. While stringent correction for relatedness reduces false positives, it may also decrease the power to detect MTAs that are correlated with population structure. To account for this, two models were implemented (i) a mixed linear model with a kinship term to account for structuring, and (ii) a relaxed linear model with the first three principal components (PCs) of relatedness included as fixed effects, to avoid overcorrection. This dual approach allowed us to assess the robustness of detected associations across models with different population structure representations. For the mixed linear model, MTAs within 250 kb of *a priori* carotenoid biosynthesis genes were considered, based on the reported LD decay of 175 – 600 kb in the SAP (Morris et al., 2013), while for the relaxed linear model, a conservative approach was taken and only MTAs within the *a priori* carotenoid biosynthesis genes were considered.

### Collections of new phenotypes, calculation of BLUPs and heritability estimates

New phenotypic data was generated through HPLC analysis of carotenoids as described in (Cruet-Burgos et al., 2026). Briefly, carotenoids were extracted from 2g samples of ground sorghum grain using a modified solid-phase extraction method (Irakli et al., 2011). An aliquot of the extract was injected through a C30 column (150 x 2 mm I.D. S-3 µm; YMC American, Inc.) at 35 °C for the carotenoid separation, and individual carotenoids (β-carotene, lutein, and zeaxanthin) were quantified using the HPLC Flexar (PerkinElmer, United States) attached to a photodiode array detector (PDA). For each sorghum accession, three biological (three separate panicles) and two technical replicates were analyzed, and an average value was calculated.

Empirical best linear unbiased predictions (eBLUPs) were calculated across *n* = 503 accessions phenotyped in three separate data sources (Figure S4-S7). Of these phenotyped accessions, *n* = 421 samples had both phenotype and genotype data and were used in pangenome-based approaches. For pangenome-based evaluations, the set of genotyped samples included *n* = 350 genotypes used in the previously described GWAS (SAP + CAP), and an additional *n* = 71 genotypes (*n* = 18 additional genotypes from (Cruet-Burgos et al., 2023), *n* = 28 genotypes from (Cruet-Burgos et al., 2026) (Global Diverse Germplasm), and *n* = 25 unpublished genotypes selected from the CAP and other sources for their grain yellowness and predicted in the top 1% of genome estimated breeding values (GEBVs) (Cruet-Burgos et al., 2023) (Selected Yellow Germplasm). Across the three sets of material, *n* = 59 genotypes were phenotyped in more than one data source. Using raw concentration values, we calculated the best linear unbiased prediction (BLUP) of random effects for each unique accession. Here, we used a linear model: *y_ij_* = μ + *b_i_* + *c_i_* +□*_ij_* where *y_ij_* is the carotenoid phenotype, μ is the global intercept, *b_i_* □□(0, σ^2^_b_) is the random effect for Source *i* phenotypes were collected in, *c* (0, σ^2^_c_) is the random effect for Accession *j*, and □*_ij_* □□(0, σ^2^□) is the residual error. Models were fit using the lme4 (v1.1-37) package in R (Bates et al., 2015). To make the value biologically relevant, we then added the conditional mean to estimate empirical BLUPs (eBLUPs), which are the phenotype values used in the text for pangenome-based approaches.

Broad-sense heritability (H²) was estimated from eBLUPs across all phenotyped individuals using the heritable (v0.1.0) package in R (Kar et al., 2026). We applied the “Standard” option, which calculates H² as the ratio of genotypic to phenotypic variance following the classical approach of (Falconer & Mackay, 1996).

### Sequencing of selected yellow grain germplasm

Accession of the ‘Selected Yellow Germplasm’ (refer to preceding section) were grown in a greenhouse under standard conditions for two weeks. Leaf tissue (approx 2 g) was collected in duplicate for each accession onto a 96-deepwell plate, flash frozen in liquid nitrogen, and shipped under dry-ice conditions to HudsonAlpha Genome Sequencing Center for sequencing. Sequencing libraries were constructed using an Illumina TruSeq DNA PCR-free library kit (Catalog #20015963) using standard protocols. Libraries were sequenced on an Illumina NovaSeq X instrument using paired ends and a read length of 150bp.

### Pangenome-based genotyping approach

The pangenome-based genotyping methodology extracts ancestry-informative pangenome-derived markers (*k*-mers) from reference assemblies for genomic regions of interest (ROIs) and counts exact matches of these markers in short-read libraries sequenced with whole-genome re-sequencing (VanGessel et al., 2025). Here we used 64-mers (*k*-mers 64bp in length) that were globally single-copy within individual pangenome reference members and absent in at least one reference member. Once all genome-wide informative markers were compiled, individual genotype hashes were generated per re-sequenced genotype (*n* = 1,124), stored as frequencies per marker for any given library.

Next, individual reference hashes were combined into a single hash, and flags were updated for only single copy 64-mers among all considered references for downstream use (e.g., a single-copy 64-mer in *reference A* is ignored if multi-copy in *reference B*). Then, genotyping hashes for the *n* = 1,124 short-read libraries sequenced with whole-genome re-sequencing were generated, storing the counts of all informative genotyping 64-mers that occur in a given library.

To isolate 64-mers that were useful for genotyping a specific region of interest (ROI), we used a graph-based method for defining the bounds of ROI syntenic sequence across reference assemblies. Here, each ROI is initially defined using -/+ 500bp the start and stop coordinates of a candidate gene in BTx623v5.1. The additional sequence added to the ROI bounds was for the inclusion of putative regulatory flanking sequences. Next, the bounds of syntenic anchor nodes (≥ 150bp in length) flanking the start and stop coordinates of the ROI are defined in each pangenome member using GraTools (v1.1.0) (Ravel et al., 2025) and extracted using bedtools (v2.31.1) (Quinlan & Hall, 2010) (Table S2). The GraTools-defined syntenic sequence for each ROI across pangenome members were *k-*merized (64-mers) and intersected with the re-sequenced short-read genotyping counts, generating raw counts of the frequency of informative 64-mers within an ROI and for each re-sequenced library. For each re-sequenced library in the raw counts matrix, we normalized raw counts by the total number of reads in a library, and set the following thresholds for generating a hard called *k-*mer presence-absence matrix: normalized counts of 0 were considered absent (0), normalized counts > 0 and < 3 were considered ambiguous (NA), normalized counts ≥ 3 were considered present (1).

This pangenome-based genotyping methodology for the generation of hard called *k-*mer presence-absence matrices was applied to 18 of the 20 biosynthesis candidate genes and the resulting number of 64-mers generated for use in genotyping each ROI is reported in Table S3. Two ROIs were excluded: *CYP97A3*;Sobic.004G346000 lacked a downstream syntenic anchor node for defining reference-specific sequence bounds and *PDS*; Sobic.006G232600 was a lower priority non-syntenic ortholog.

### Pangenome-based evaluation of regions of interest

All pangenomic analyses in this study were performed using the sorghum pangenomic resources described by (Morris et al., 2026). GENESPACE (v.1.3.1) (Lovell et al., 2022) was used to determine geneIDs of syntenic orthologs for the carotenoid biosynthesis candidate genes across pangenome members. To evaluate predicted transcript isoform diversity of the *ZEP* gene across pangenome members, we extracted transcript annotation information from annotation files for each pangenome reference assembly, available on Phytozome (http://www.phytozome.net) (Goodstein et al., 2012; Morris et al., 2026), and visualized the transcript isoforms using custom scripts. To visualize presence-absence variation within each ROI, an ROI-specific variant call file (VCF) was derived from the pangenome graph via vg toolkit (v1.61.0) (Garrison et al., 2018; Liao et al., 2023) using the syntenic bounds determined above (Table S2) and PAVplotR (https://github.com/avril-m-harder/PAVplotR) was used to calculate an expanded coordinate system for variants in each interval (Harder, 2026). The expanded coordinate system allowed for consideration of insertions relative to the primary reference (BTx623) and for calculation of proportional sequence presence in 100bp bins.

### Detecting associations with pangenome-derived markers

To assess the significance for presence/absence of individual 64-mers association to variation in carotenoid traits (β-carotene, lutein, zeaxanthin), we extracted the false discovery rate (FDR) adjusted *p*-value from *t*-tests. For *t*-tests, we required a minimum of two marker call groups (presence, absence, ambiguous) with ≥ five samples per group. For ROIs where we explored the relative structure of these 64-mer trait associations in a genomic interval, we determined the 64-mer’s relative position in multiple sequence alignment (MSA) space. MSAs were generated with mafft (v7.525) (Katoh & Standley, 2013) under default settings and the ungapped position of 64-mers were determined within the ROI of the MSA using custom scripts.

### Using machine learning to detect associations across candidate genes in the carotenoid biosynthetic pathway

We hypothesized that a machine learning (ML) framework would provide a valuable approach for modeling carotenoid trait-genotype relationships across many markers at once and for the prioritization across our list of *a priori* carotenoid biosynthesis candidate genes. To test this, we constructed ML models for the three carotenoid traits (β-carotene, lutein, zeaxanthin) separately and using a set of pangenome-derived markers across *n* = 18 *a priori* candidate biosynthesis genes. Models were built with eXtreme Gradient Boosting (XGBoost), a ML method which operates based on gradient boosted decision trees (T. Chen & Guestrin, 2016; W. Li et al., 2019; Montomoli et al., 2021). To reduce the number of features included in model construction, for each candidate, we removed highly correlated (r >= 0.9) pangenome-derived markers (Table S3, pangenome-derived markers; total *n* = 294,906, non-redundant *n* = 11,700). Using the set of non-redundant pangenome-derived markers and the same set of phenotypes used in the local pangenome-based associations (*n* = 421), data was split into training and testing (70% training, 30% testing) and the training set data (*n* = 294) was used in model tuning and validation.

Gradient boosting models were trained using XGBoost (v1.7.11.1) in R with a regression objective function (reg:squarederror) to predict quantitative values of the three carotenoid traits. Hyperparameter tuning was performed using a grid search with a 5-fold cross-validation on the training set. The search space was testing using the xgboost::xgb.cv function and included the following parameters: learning rate (eta = c(0.01, 0.05, 0.1)), tree depth (max_depth = c(3, 4, 5)), row sampling (subsample = c(0.7, 0.8, 0.9)), and column sampling (colsample_bytree = c(0.1, 0.2, 0.3)). The optimal parameter set was selected based on the lowest cross-validation root mean squared error (RMSE) across all three trait models, so that a single set of parameters were used for the final trait-specific models. The selected parameters, eta = 0.05, max_depth = 5, subsample = 0.9, colsample_bytree = 0.1, were used to train the final models on the training set, and the number of boosting rounds was determined individually for each trait based on the cross-validation stopping point.

Final models were evaluated on the held-out testing set using three metrics, RMSE, mean absolute error (MAE), and coefficient of determination (R^2^). Feature importance was extracted from each trained model using the XGBoost gain metric, which measures the contribution of each feature to reducing loss during tree splits. Features were aggregated by genomic region of interest (ROI) by summing gain across pangenome-derived markers belonging to the same ROI. A fixed random seed was used throughout all stochastic operations (train-test splitting, cross-validation folds) to ensure reproducibility

### Author contributions

LB conducted the investigation, formal analysis of carotenoid data , writing - original draft, and review and editing. CMM conducted the investigation, formal analysis of pangenomic data (bioinformatics), visualization, writing - original draft, and review and editing. AMH contributed to the methodology, software, visualization, validation, and writing - review and editing. CCB contributed to writing - review and editing. ALH contributed to the methodology, software, validation, data curation, and writing - review and editing. DF contributed to the methodology, software, and data curation. JG, AS, and ST contributed to data acquisition and data curation (sequencing). GPM conceptualized the study and contributed to writing - review and editing, supervision, and funding acquisition. JTL performed the data curation, and contributed to writing - review and editing, supervision, and funding acquisition. DHR conceptualized the study and contributed to writing - review and editing, supervision, and funding acquisition. All authors read and approved the manuscript.

## Data availability

The sorghum pangenomic resources used in this study are described by (Morris et al., 2026) and are available through Phytozome under the SorghumPan project (https://phytozome-next.jgi.doe.gov/sorghumpan/). All raw sequencing reads for ’Selected Yellow Germplasm’ accessions have been deposited in the NCBI SRA database under BioProject accession PRJNA1531092. Previous carotenoid concentration data is reported in (Cruet-Burgos et al., 2023) and (Cruet-Burgos et al., 2026). New carotenoid concentration data and additional data/figures supporting the findings of this study are available in the Supporting Data Files included with this article.

