## Supplemental File 2 Figures S1-S16 for "Pangenomics and machine learning reveal genetic variation to optimize carotenoids in sorghum grain"

**SUPPLEMENTARY FILE 2**

**FIGURES S1-S16**

**Pangenomics and machine learning reveal genetic variation to optimize carotenoids in sorghum grain**

Linly Banda ^1^**^†^**, Chloee M. McLaughlin ^1, 2^**^†^**, Avril M. Harder ^2^, Clara Cruet-Burgos ^3^, Adam L. Healey ^2^, Dave Flowers ^2^, Shannon Talley ^2^, Ada Stewart ^2^, Jane Grimwood ^2^, Geoffrey P. Morris ^3^, John T. Lovell *^2^ and Davina H. Rhodes *^1^

^1^ Department of Horticulture and Landscape Architecture, Colorado State University, Fort Collins, CO 80523, USA

^2^ Genome Sequencing Center, HudsonAlpha Institute for Biotechnology, Huntsville, AL 35806, USA

^3^ Department of Soil and Crop Science, Colorado State University, Fort Collins, CO 80523, USA

**^†^** equal contribution

*Corresponding authors:

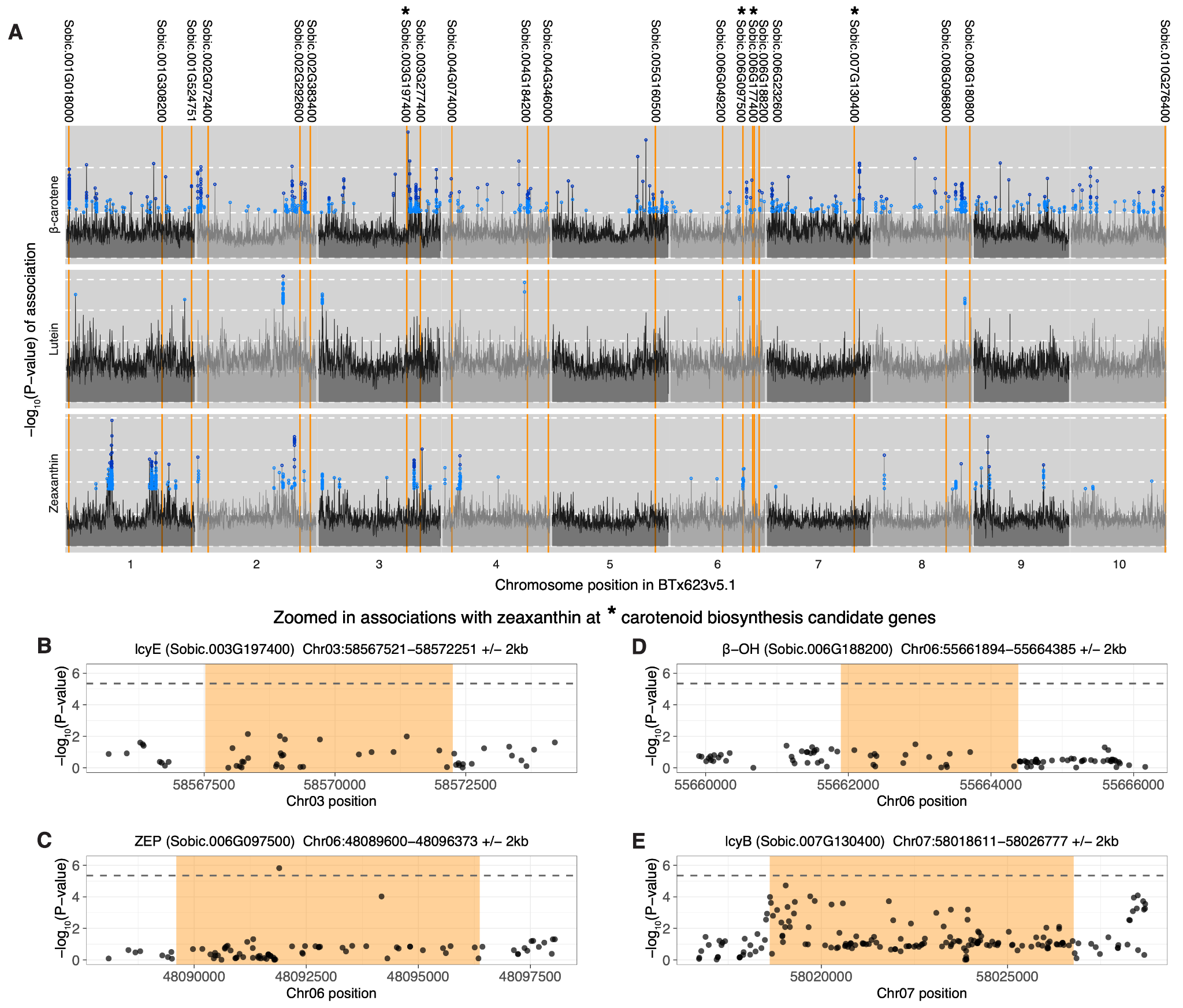

### Figure S1

**Genome-wide association of carotenoid traits using a model including a population structure covariate.** A) Genome-wide association with carotenoid traits (top: β-carotene, middle: lutein, bottom: zeaxanthin) using a linear mixed model (LMM), accounting for a relatedness matrix. For all traits, the maximum -log_10_(*P*-value) association within non-overlapping 100kb intervals is plotted genome-wide (black/gray lines). All significant associations are plotted as colored points (dark blue: FDR *P*-value < 0.005, light blue: FDR *P*-value < 0.05). Orange vertical lines are the genomic position of biosynthesis candidate genes, and the gene model is given above plots. Gene models with an asterisks (*) are plot in zoomed in association panels B) *LcyE* (Sobic.003G197400) C) *ZEP* (Sobic.006G097500) D) *β*-*OH* (Sobic.006G188200) E) *LcyB* (Sobic.007G130400). Panels B-E are zoomed in associations with the zeaxanthin trait. Points plot include +/- 2kb of the gene model, the bounds of the gene model are highlighted in orange, and the dashed line is FDR *P*-value < 0.05. Zeaxanthin association in panel A uses the same data as in Figure 1A.

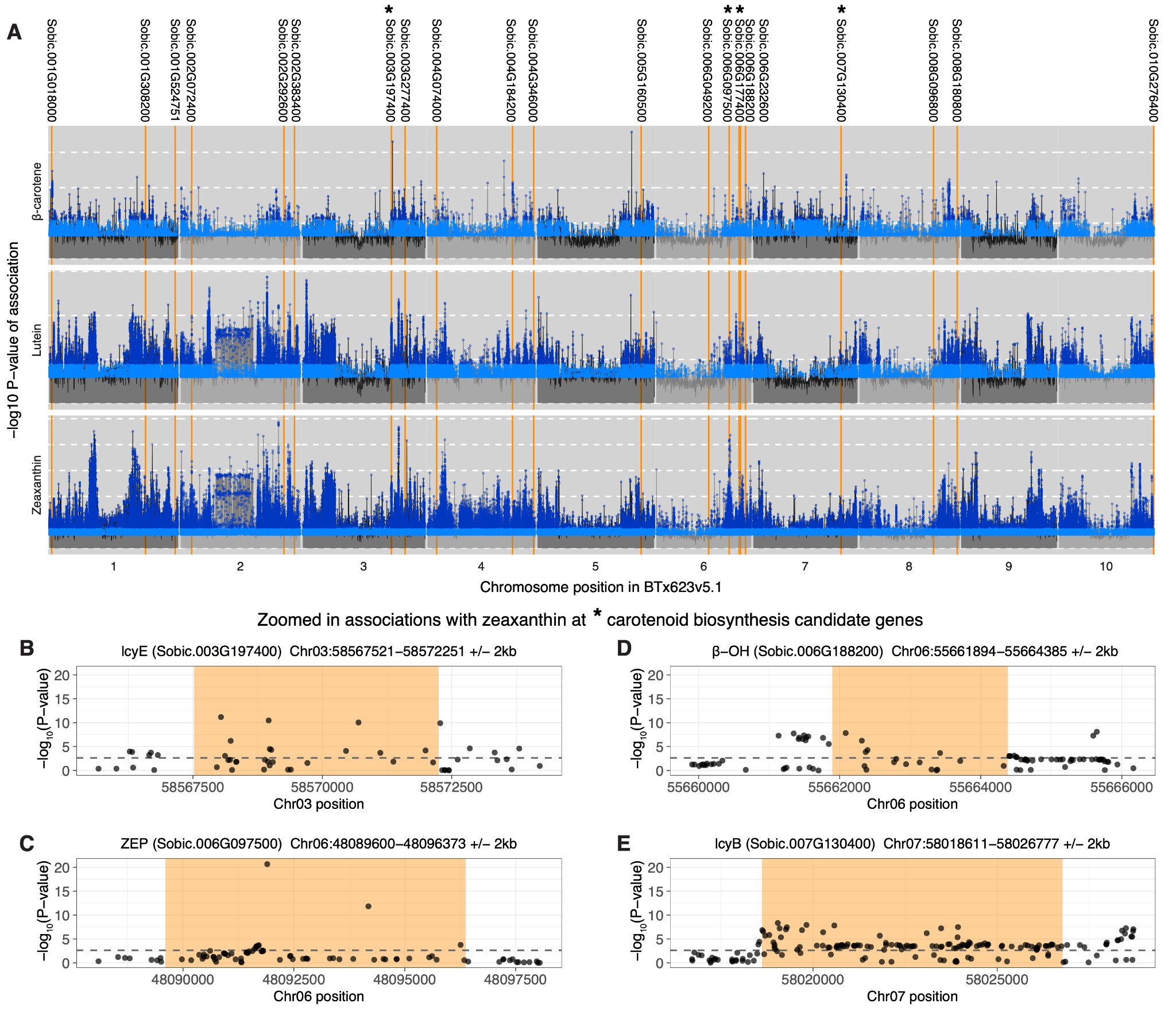

### Figure S2

**Genome-wide association of carotenoid traits using a model with a more relaxed population structure correction.** A) Genome-wide association with carotenoid traits (top: β-carotene, middle: lutein, bottom: zeaxanthin) using a linear model, accounting for the first 3 principal components of a relatedness as fixed effects. For all traits, the maximum -log_10_(*P*-value) association within non-overlapping 100kb intervals is plotted genome-wide (black/gray lines). All significant associations are plotted as colored points (dark blue: FDR *P*-value < 0.005, light blue: FDR *P*-value < 0.05). Orange vertical lines are the genomic position of biosynthesis candidate genes, and the gene model is given above plots. Gene models with an asterisks (*) are plot in zoomed in association panels B) *LcyE* (Sobic.003G197400) C) *ZEP* (Sobic.006G097500) D) *β*-*OH* (Sobic.006G188200) E) *LcyB* (Sobic.007G130400). Panels B-E are zoomed in associations with the zeaxanthin trait. Points plot include +/- 2kb of the gene model, the bounds of the gene model are highlighted in orange, and the dashed line is FDR *P*-value < 0.05.

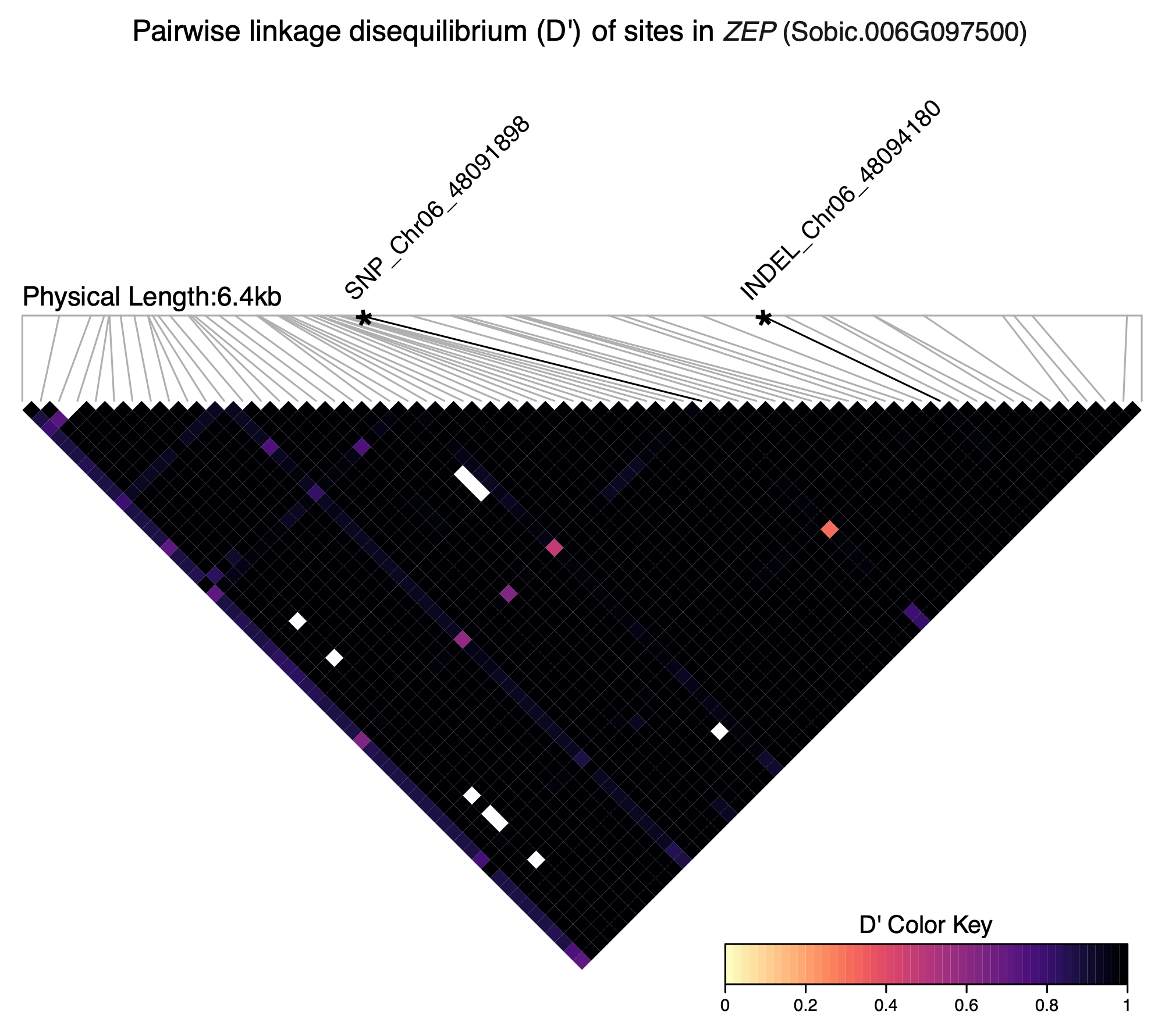

### Figure S3

**Linkage disequilibrium heatmap of the genomic interval containing *ZEP*.** Pairwise linkage disequilibrium heatmap for variants of *ZEP* (Sobic.006G097500) Chr06:48089600-48096373 called against the linear BTx623v5.1 coordinate system. Asterisks and black lines are annotated for the variants SNP_Chr06_48091898 and INDEL_Chr06_4809418.

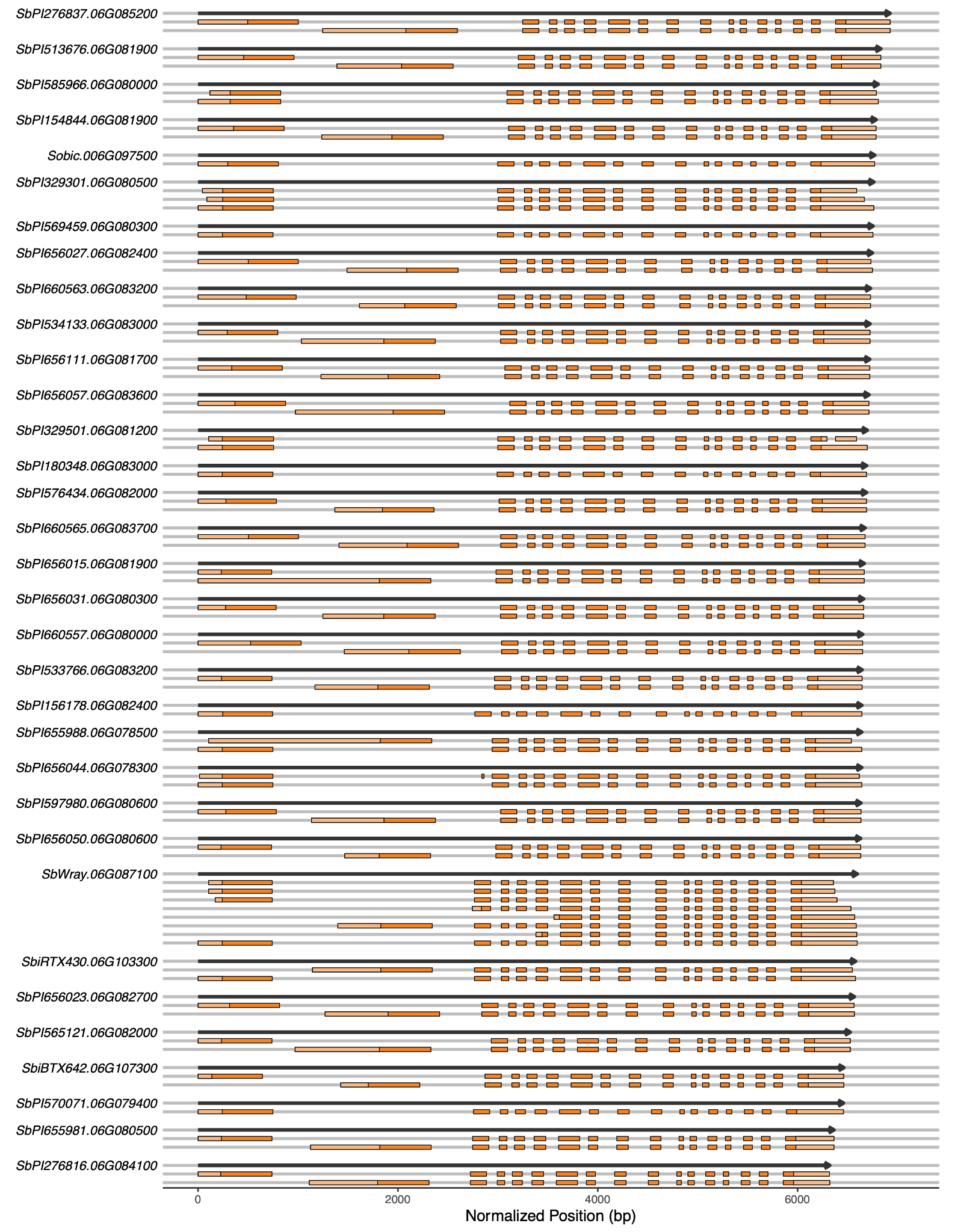

### Figure S4

**Gene structure and isoform diversity of *ZEP* (Sobic.006G097500) across *n* = 33 sorghum pangenome reference members.** For each pangenome member, the full-length gene model is drawn in black, with transcript isoforms that are reported in each reference’s annotation file drawn below. Features of isoforms are color-coded, the light orange boxes denote 5' (left-most) and 3' (right-most) untranslated regions (UTRs), dark orange boxes indicate exons. All gene models and transcript isoforms are plotted to a normalized coordinate system, based on the relative length of each gene model. Gene models and corresponding transcript isoforms are plotted in descending order of the full length gene model. Transcripts SbiRTx430.06G103300 and Sobic.006G097500 are the same as in Figure 2. Order of pangenome reference member (PI number) from top to bottom - PI276837, PI513676, PI585966, PI154844, PI564163 (BTx623), PI329301, PI569459, PI 656027, PI660563, PI534133, PI656111, PI656057, PI329501, PI180348, PI576434, PI660565, PI656015, PI656031, PI660557, PI533766, PI156178, PI655988, PI656044, PI597980, PI656050, PI653616 (Wray), PI655996 (RTx430), PI656023, PI565121, PI656029 (BTx642), PI570071, PI655981, PI276816.

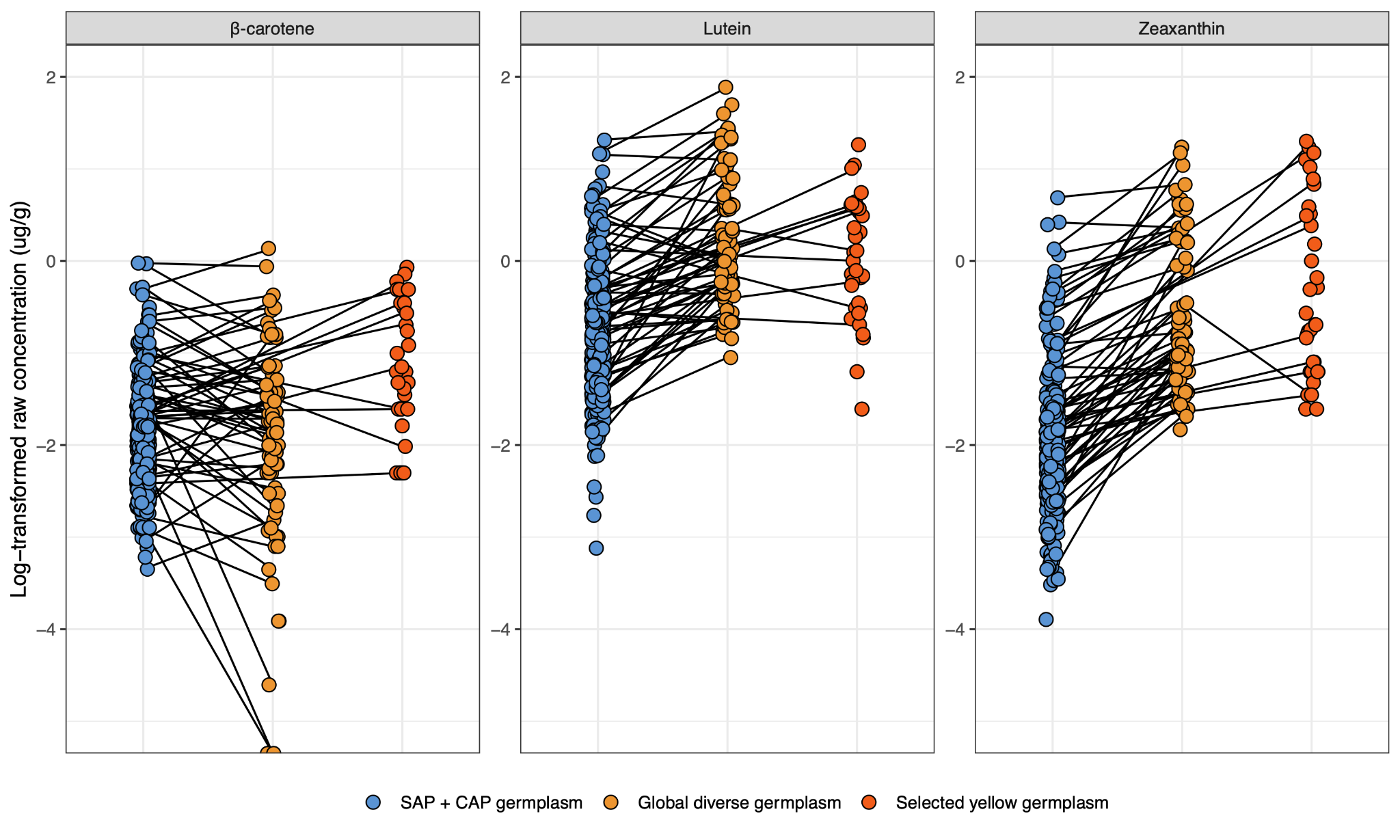

### Figure S5

**Raw log-transformed concentrations of carotenoid traits used in the calculation of eBLUPs.** For each trait (β-carotene, lutein, zeaxanthin) points are an accession’s average phenotype value collected in a source: SAP + CAP germplasm (Cruet-Burgos 2023), global diverse germplasm (Cruet-Burgos, Banda et al. 2026), selected yellow germplasm (novel and previously unexplored germplasm) and lines connect accessions that were phenotyped across multiple sources.

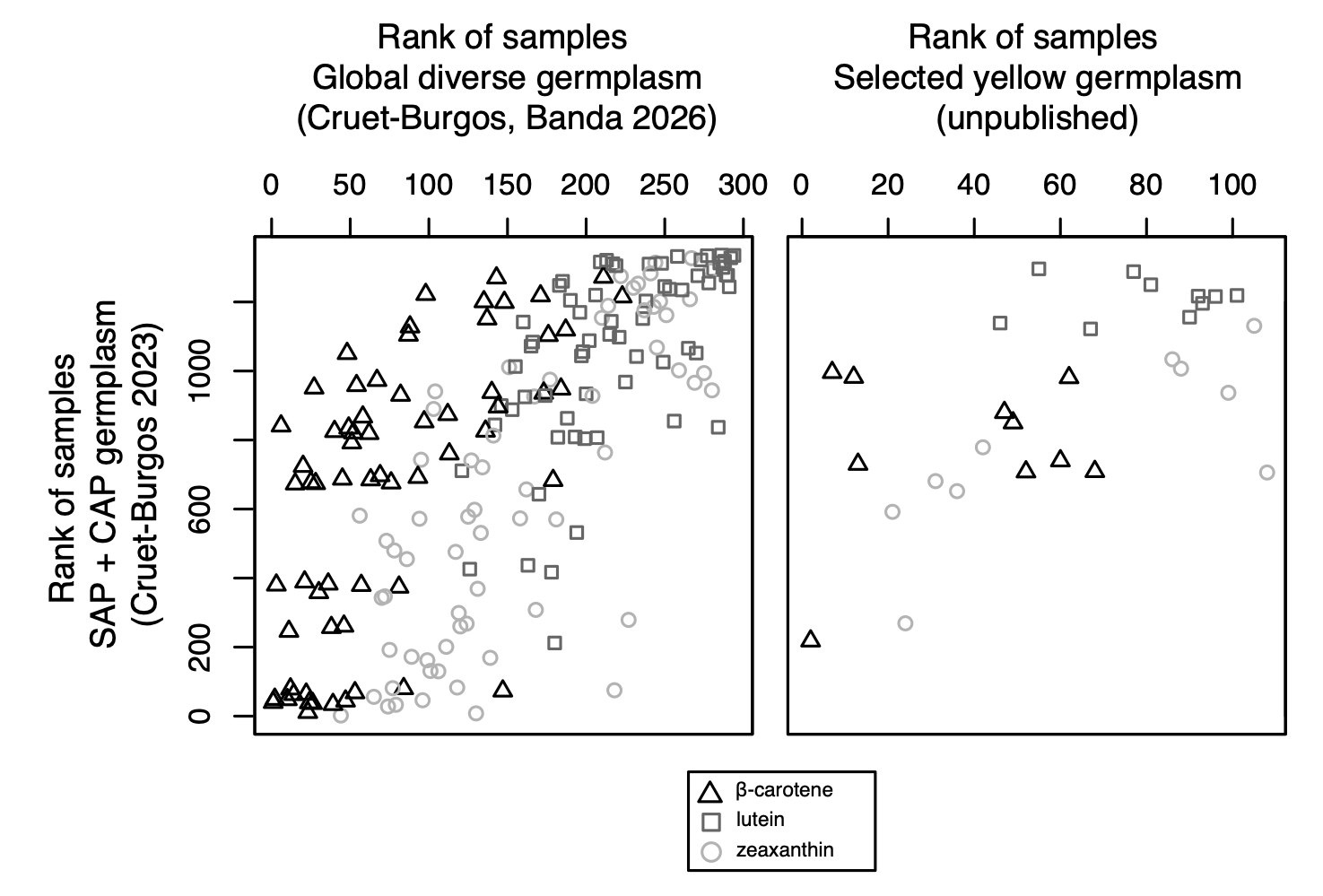

### Figure S6

**Rank of raw trait values for germplasm phenotyped across multiple sources.** For each trait (β-carotene, triangle; lutein, square; zeaxanthin, circle) points are the raw ranked trait values of accessions phenotyped in multiple sources.

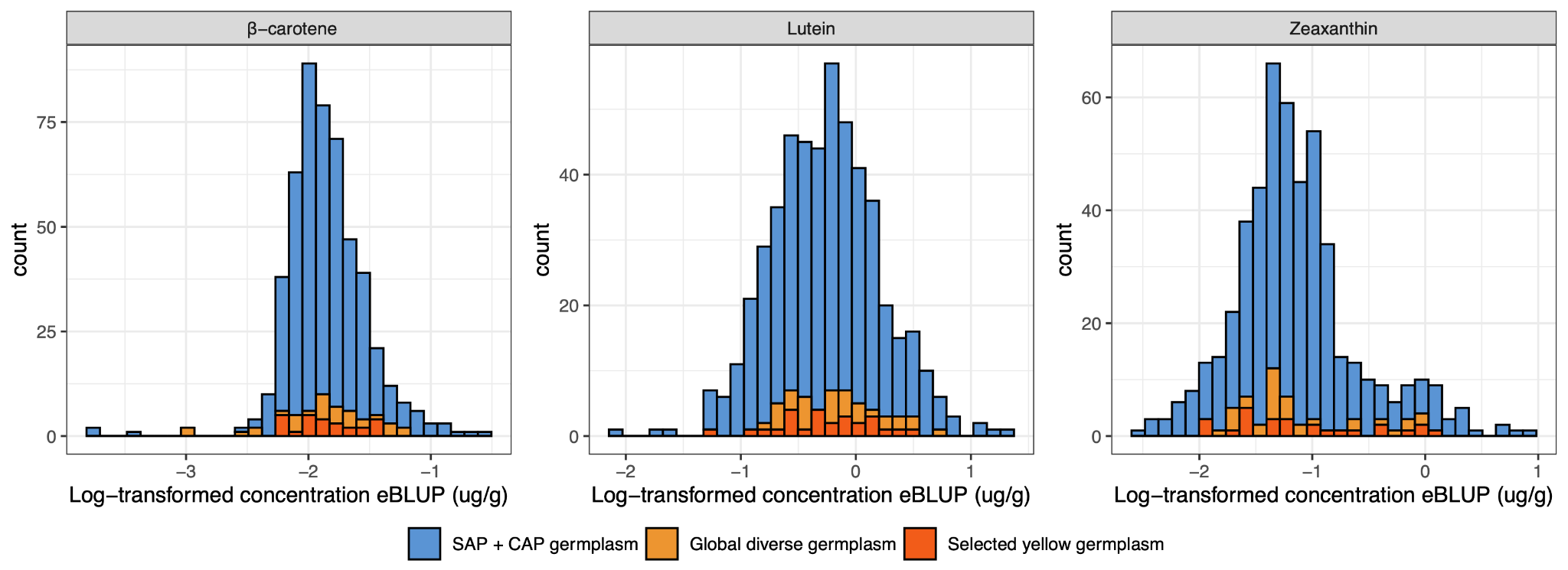

### Figure S7

**Distribution of eBLUPs by source for carotenoid traits.** Histograms showing the range of empirical best linear unbiased predictors (eBLUPs) for β-carotene, lutein, and zeaxanthin across *n* = 503 phenotyped unique accessions. Bars are colored by phenotyping source: SAP + CAP germplasm (includes accessions phenotyped in Cruet-Burgos et al. 2023 and replicated in other sources), global diverse germplasm (novel material published in Cruet-Burgos, Banda et al. 2026), and selected yellow germplasm (novel and previously unexplored germplasm only)

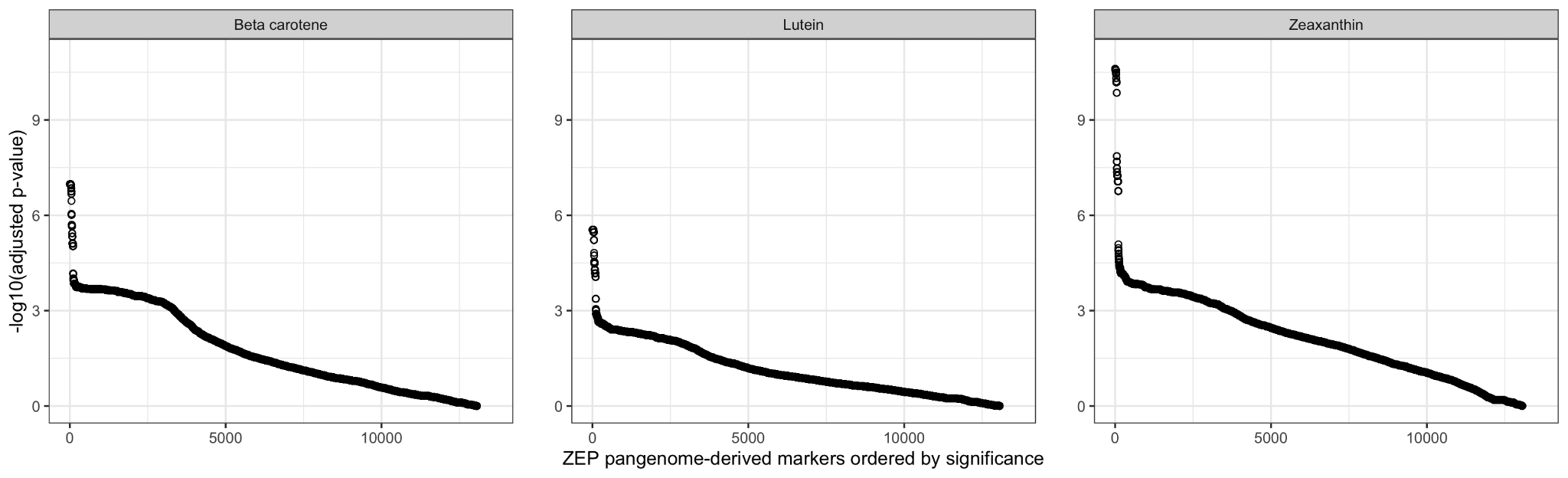

### Figure S8

**Significance of *ZEP* (Sobic.006G097500) genotyping pangenome-derived markers.** Each point is a pangenome-derived marker of the *ZEP* genomic region (*n* = 13,550 markers in total) and ordered by the significance of -log_10_(adjusted *P*-value) for association to the three carotenoid traits. Points of the zeaxanthin pane are the same as in Figure 3D, and here are ordered by significance rather than in a shared common coordinate system.

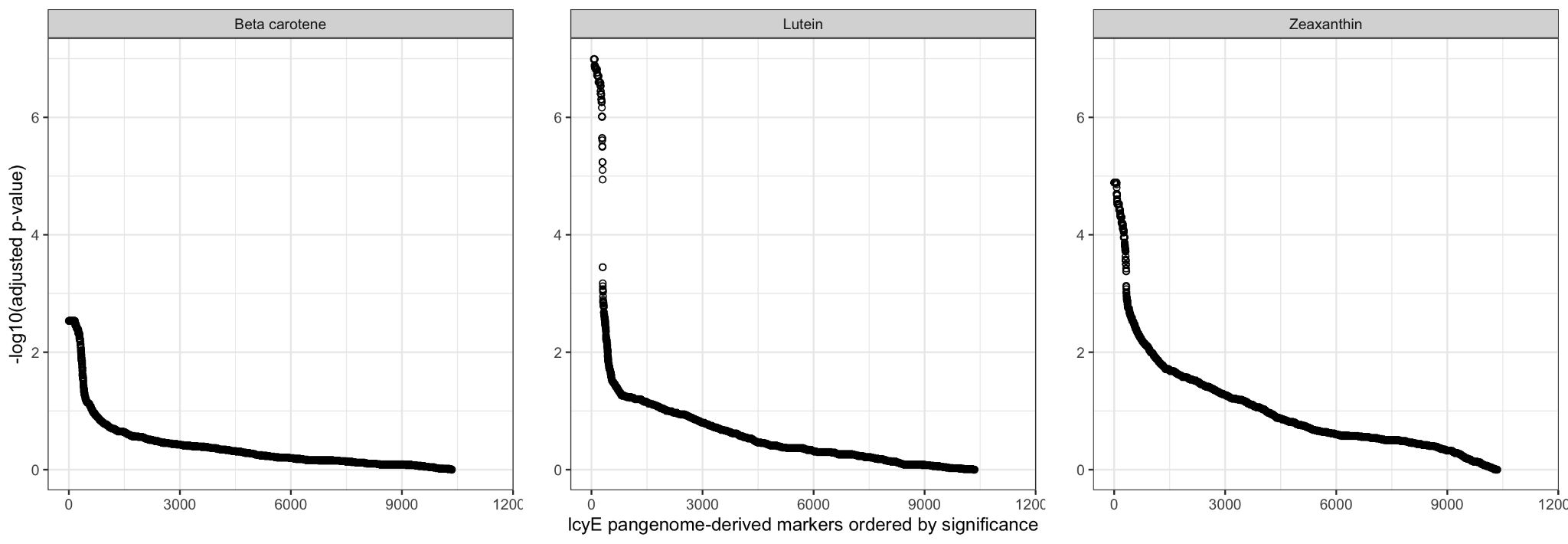

### Figure S9

**Significance of *LcyE* (Sobic.003G197400) genotyping pangenome-derived markers.** Each point is a pangenome-derived marker of the *LcyE* genomic region (*n* = 11,445 markers in total) and ordered by the significance of -log_10_(adjusted *P*-value) for association to the three carotenoid traits. Points of the lutein pane are the same as in Figure S10, and here are ordered by significance rather than in a shared common coordinate system.

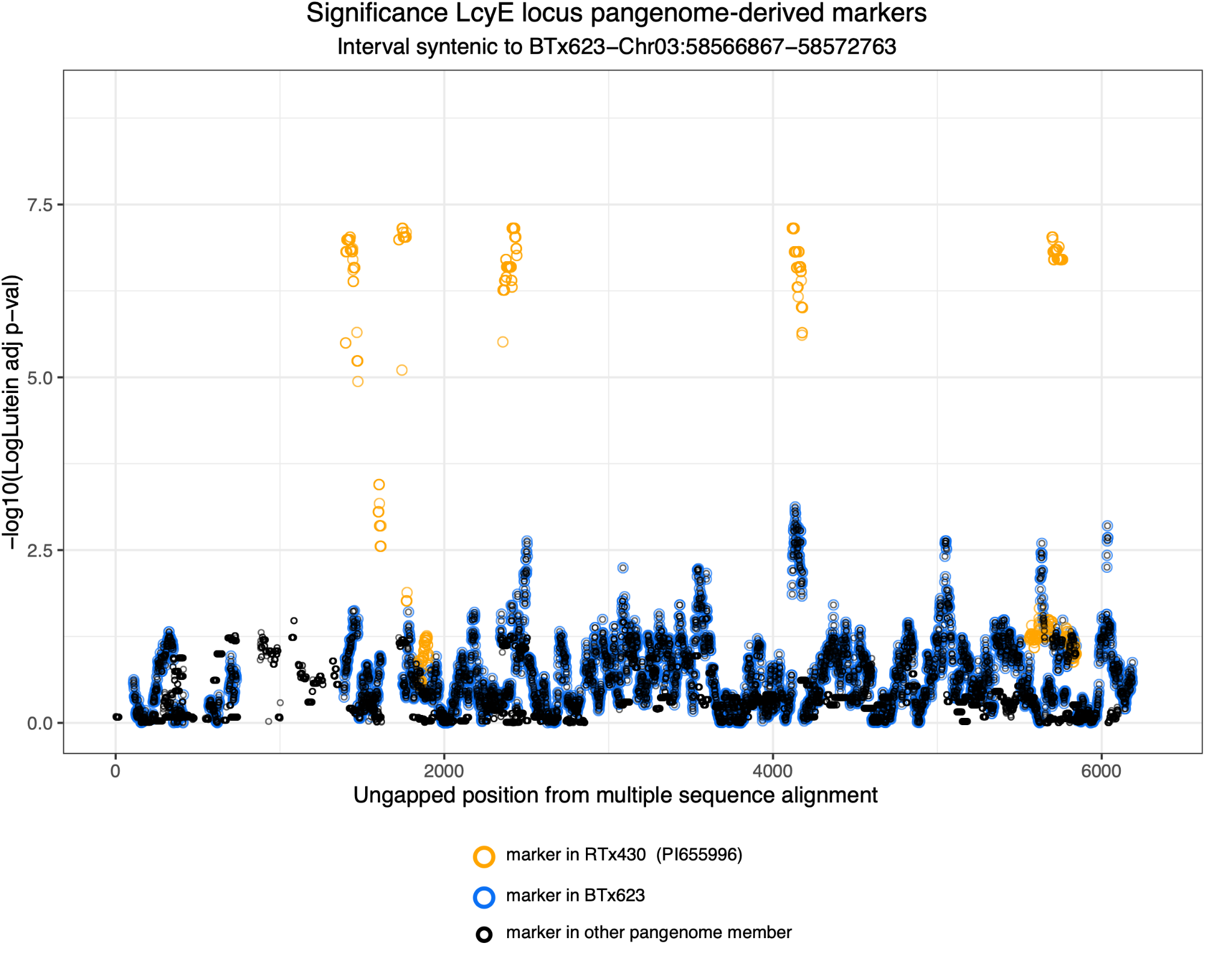

### Figure S10

**Association of *LcyE* (Sobic.003G197400) pangenome-derived markers with variation in lutein content plotted in a shared coordinate system.** -log_10_(adjusted *P*-value) of association with lutein using the pangenome-derived markers (64-mers). Orange points are pangenome-derived markers found in RTx430, blue points are pangenome-derived markers found in BTx623v5.1, and black points are pangenome-derived markers not represented in RTx430 or BTx623v5.1, but present in the other *n* = 31 pangenome members.

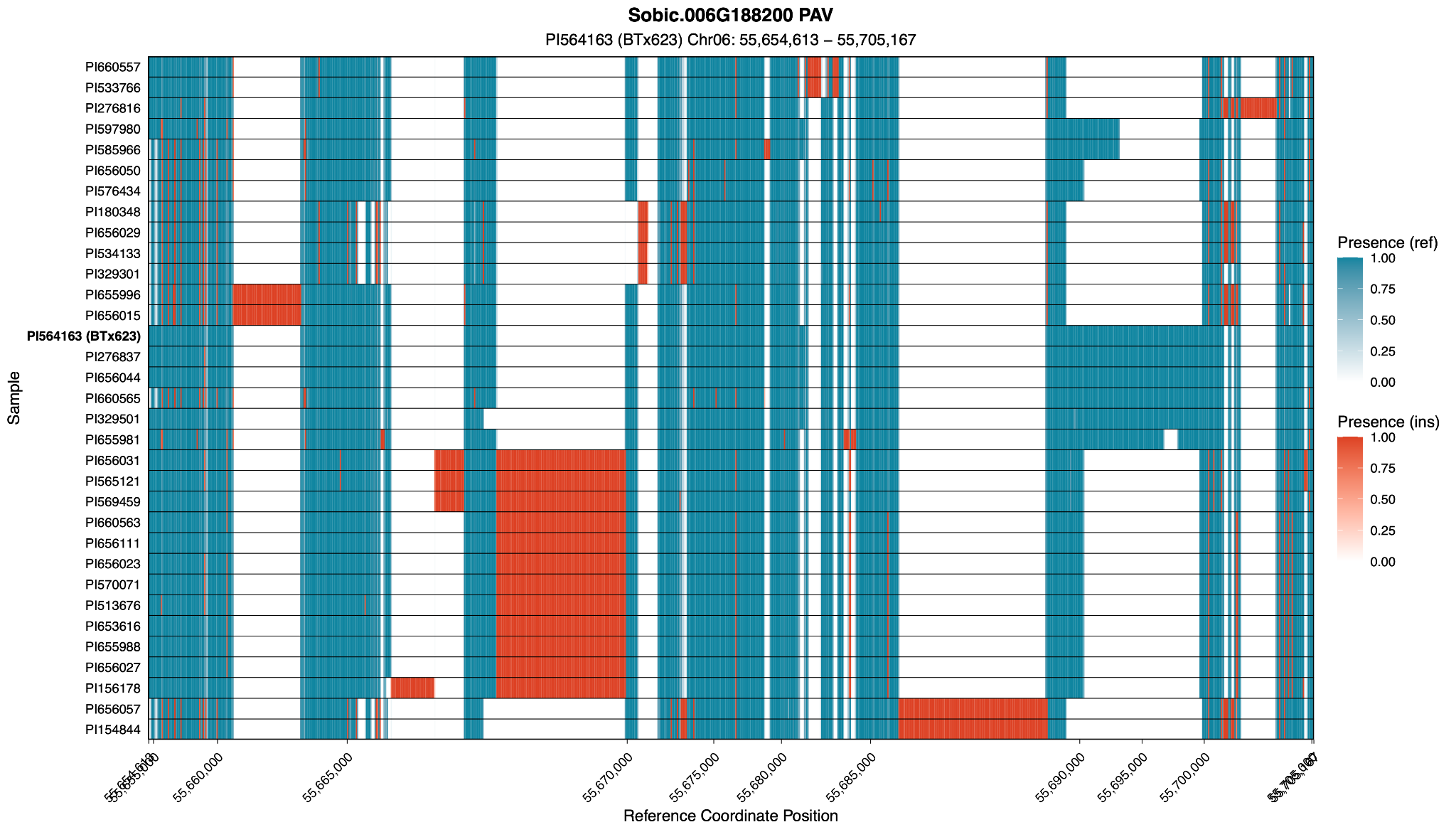

### Figure S11

**Presence-absence variation at *β-OH* (Sobic.006G188200) genomic region across 33 sorghum pangenome reference members.** Sequence presence-absence variation in *n* = 33 sorghum pangenome reference members (rows) for the genomic region containing *β-OH* (Sobic.006G188200) syntenic to BTx623v5.1 Chr06: 55,654,613 − 55,705,167. Each line is a 100bp bin colored blue by the proportion of present sequence of the primary reference BTx623 (Presence (ref)) or red by the proportion of inserted sequence relative to the BTx623 (Presence (ins)).

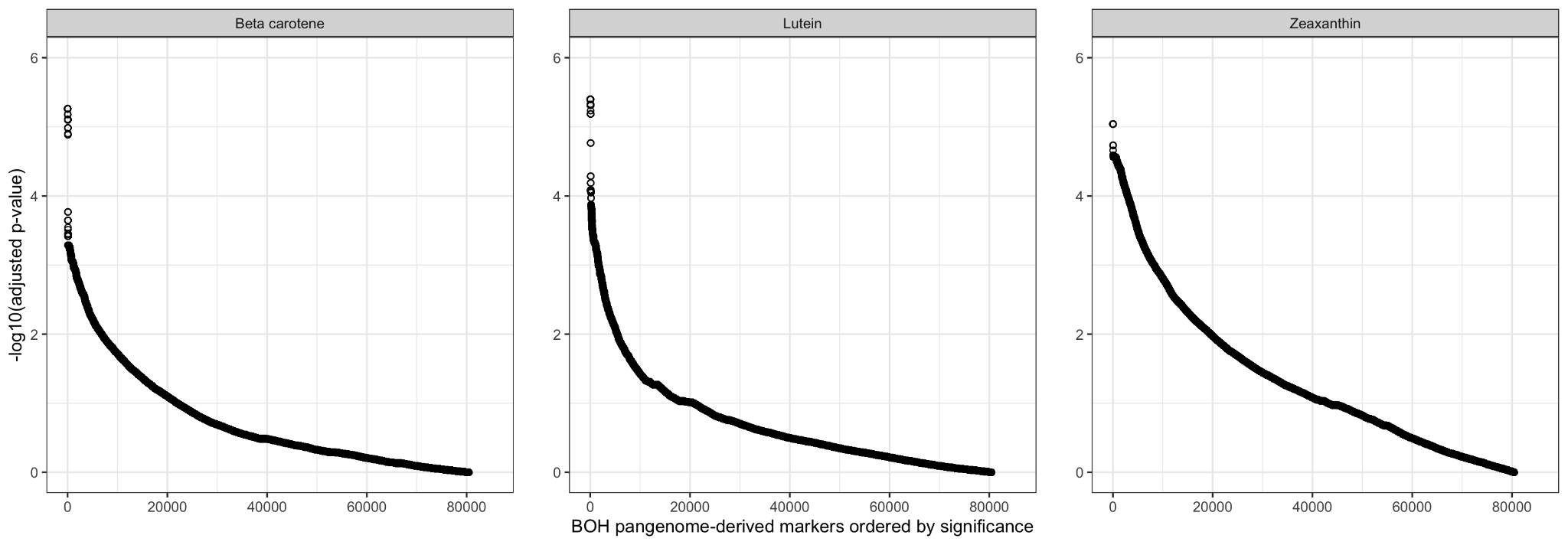

### Figure S12

**Significance of *β-OH* (Sobic.006G188200) genotyping pangenome-derived markers.** Each point is a pangenome-derived marker in the *β-OH* genomic region (*n* = 85,179 markers in total) and ordered by the significance of -log_10_(adjusted *P*-value) for association to the three carotenoid traits.

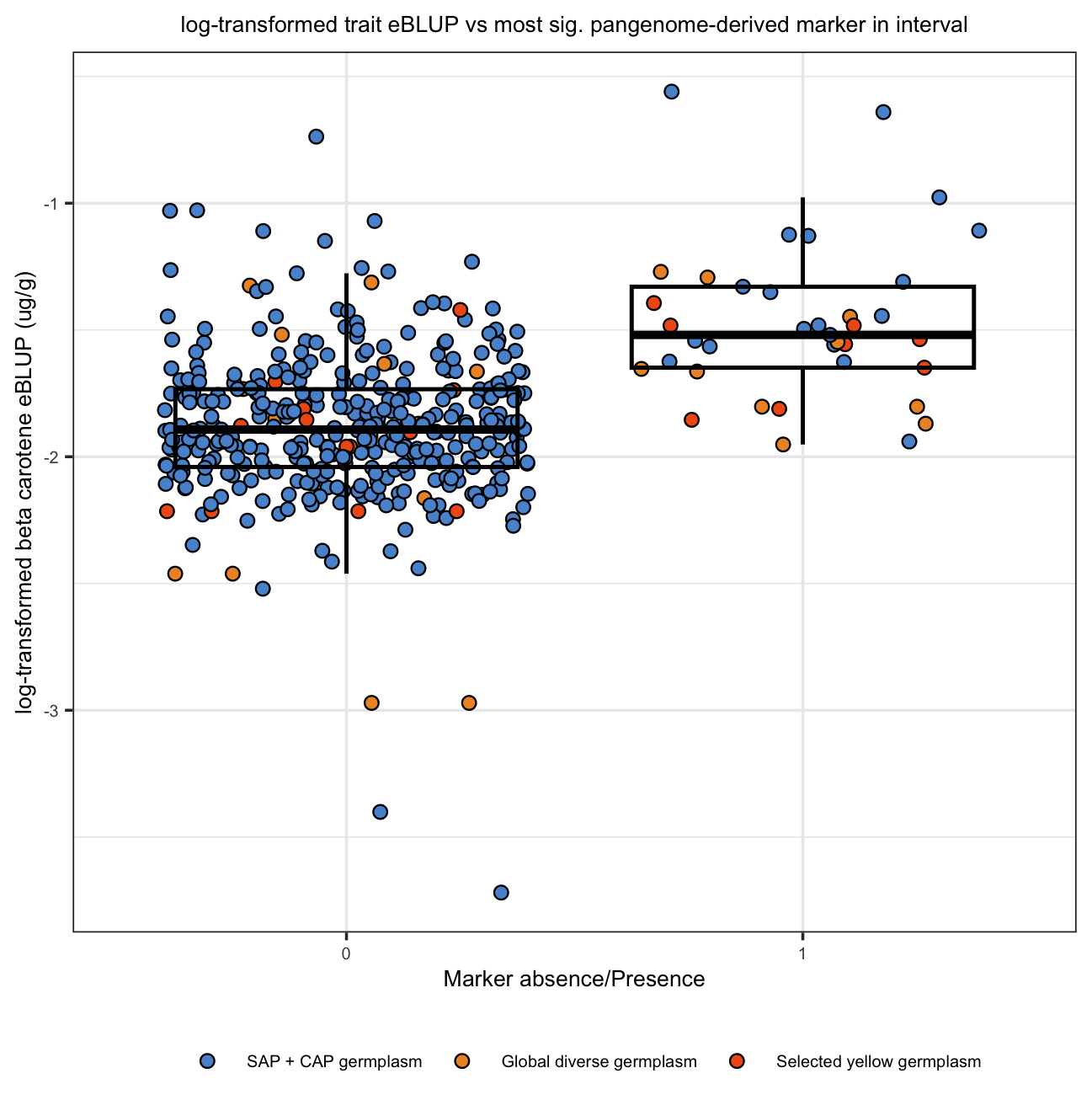

### Figure S13

**Variation in beta carotene content is associated with the most significant *β-OH* pangenome-derived marker**. Boxplots of log-transformed beta carotene concentrations across genotypes of the most significant pangenome-derived marker in the *β-OH* (Sobic.006G188200) region (absence = 0, presence = 1) for sorghum germplasm used in the pangenome-based association studies (*n* = 421). Sorghum accessions are colored by phenotyping source: SAP + CAP germplasm (includes genotypes phenotyped in Cruet-Burgos et al. 2023 and replicated in other sources), global diverse germplasm (novel material published in Cruet-Burgos, Banda et al. 2026), and selected yellow germplasm (novel and previously unexplored germplasm only). One individual with an ambiguous genotype call of the Selected yellow germplasm group was not plotted and had a phenotype value of log-transformed beta carotene concentration = -2.21.

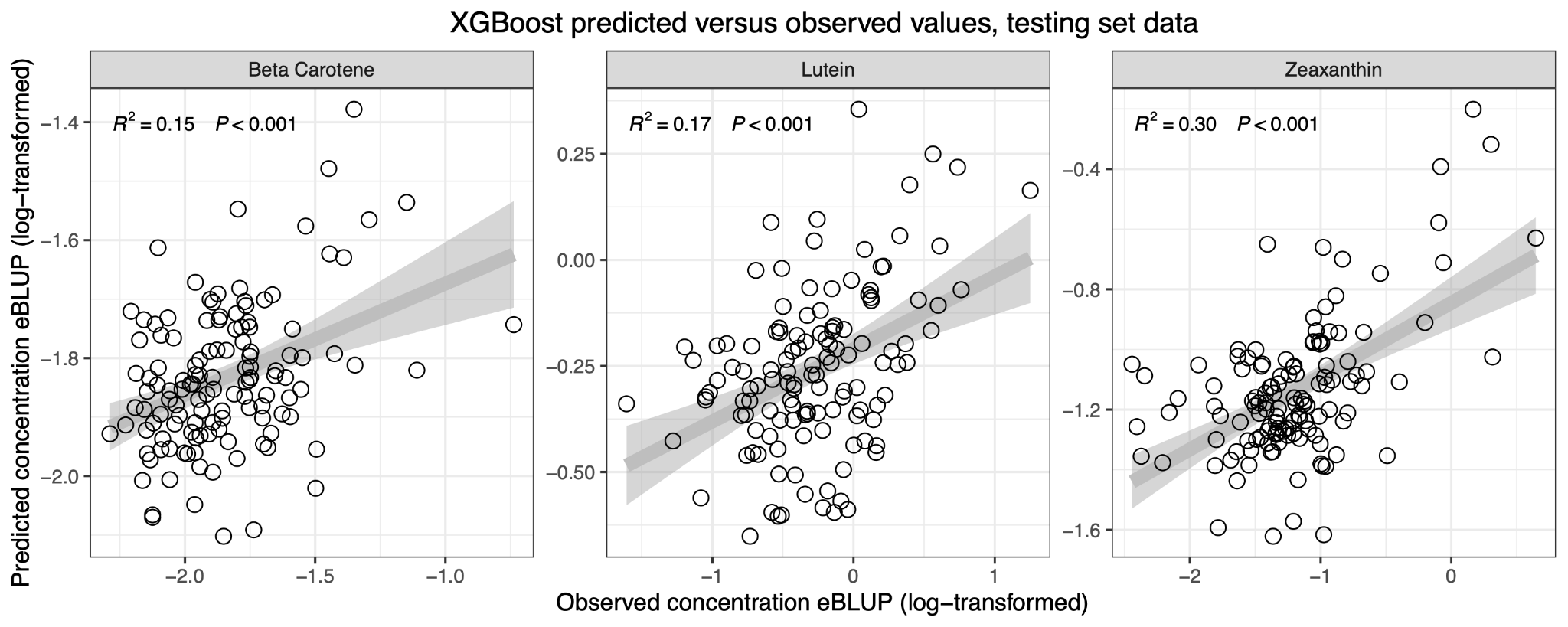

### Figure S14

**XGBoost trait predictions on testing set (hold-out) data.** For the three separate XGBoost trait models, predicted versus observed trait values for the *n* = 127 accessions of the testing (hold out) data not used to train XGBoost models. The zeaxanthin panel is identical to that shown in Figure 4B.

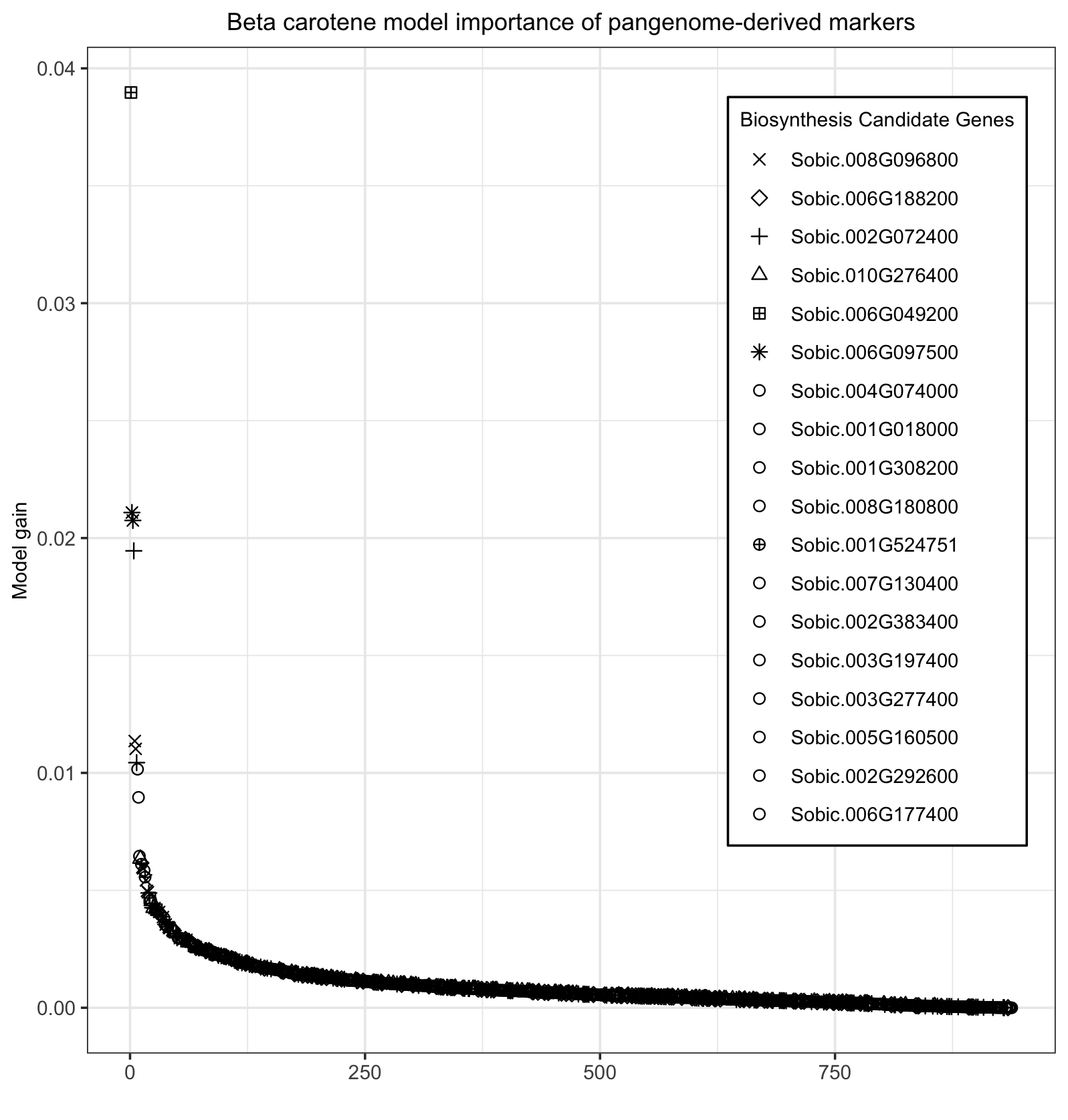

### Figure S15

**Pangenome-derived markers used in the β-carotene XGBoost model ordered by gain**. Pangenome-derived markers (*n* = 938) selected for training the XGBoost model for β-carotene prediction, ordered by gain (a measure of feature importance). Each point represents a marker and is styled (shape) according to the carotenoid biosynthesis candidate gene to which it belongs. Biosynthesis candidate genes in the legend are ordered by the aggregated gain of markers, where the first gene reported accounts for the largest summed gain. Markers of candidate genes with a distinct point shape share the same styling as in Figure 4C and Figure S16.

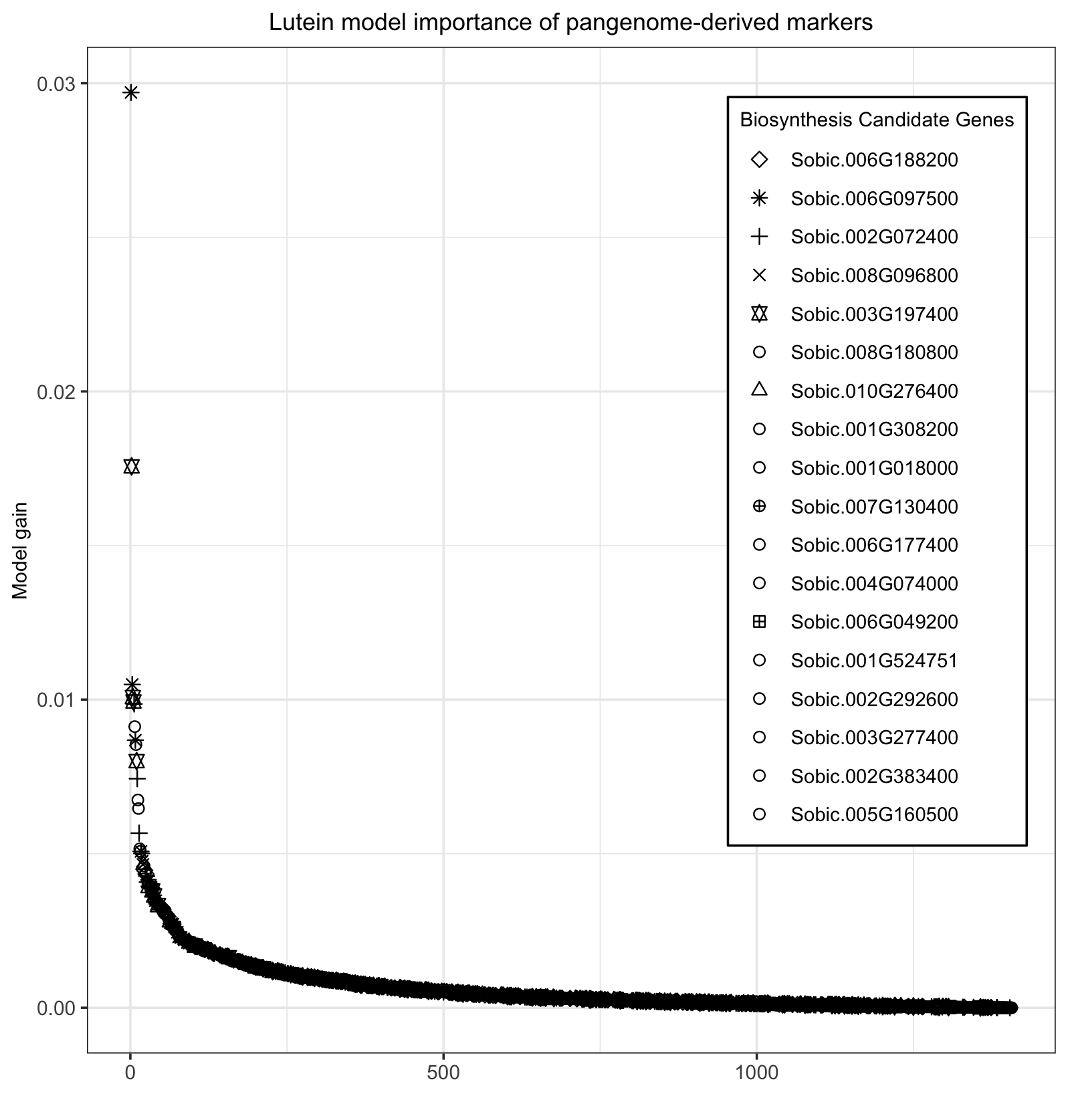

### Figure S16

**Pangenome-derived markers used in the lutein XGBoost model ordered by gain**.

Pangenome-derived markers (*n* = 1,407) selected for training the XGBoost model for lutein prediction, ordered by gain (a measure of feature importance). Each point represents a marker and is styled (shape) according to the carotenoid biosynthesis candidate gene to which it belongs. Biosynthesis candidate genes in the legend are ordered by the aggregated gain of markers, where the first gene reported accounts for the largest summed gain. Markers of candidate genes with a distinct point shape share the same styling as in Figure 4C and Figure S15.
